# Sequence Composition Dictates Condensate Miscibility

**DOI:** 10.1101/2024.11.29.626135

**Authors:** Gaofeng Pei, Xinxin Wang, Danqian Geng, Zhuo Chen, Weifan Xu, Tingting Li, Pilong Li

**Affiliations:** State Key Laboratory of Membrane Biology, Frontier Research Center for Biological Structure, School of Life Sciences, Tsinghua University; Tsinghua University-Peking University Joint Center for Life Sciences, Beijing 100084, China; Department of Biochemistry and Molecular Biology, School of Basic Medical Sciences, Peking University Health Science Center, Beijing, 100191, China; State Key Laboratory of Integrated Management of Pest Insects and Rodents, Institute of Zoology, Chinese Academy of Sciences; CAS Center for Excellence in Biotic Interactions, University of Chinese Academy of Sciences, Beijing, China

**Keywords:** Biomolecular condensate, Miscibility, Intrinsically disordered regions, Gene expression regulation

## Abstract

Numerous biomolecular condensates coexist within cells, yet the mechanisms governing their mixing and demixing remain elusive. To investigate how the amino acid composition of intrinsically disordered regions (IDRs) affects condensate coexistence, we paired and tested 28 IDRs in 378 different combinations. Our results reveal that IDRs enriched in serine or aromatic amino acids tend to form miscible condensates. Conversely, IDRs with high charge levels form immiscible condensates, even when the serine levels are artificially boosted. Additionally, phosphorylation acts as a switch, modulating condensate miscibility. We also observed that the miscibility between transcription factor condensates and Pol II condensates profoundly influences transcription. We engineered transcription factors by increasing serine content, enhancing both their miscibility with Pol II and their transcriptional activity. However, introducing high levels of charge had the opposite effect. These findings shed light on the fundamental mechanisms controlling condensate miscibility and offer insights for designing specific functional condensates.

## Introduction

Over the past decade, dozens of types of biomolecular condensates have been identified in eukaryotic cells. Considerable efforts have been devoted to studying their formation mechanisms, components, selective partitioning, and functions. However, the mechanisms that control the coexistence behavior of different condensates have yet to be adequately explored.

Biomolecular condensates are formed, in part, by phase separation driven by multivalent interactions^1^ among biomacromolecules. For proteins, these interactions can arise within modular domains^1^ or within intrinsically disordered regions (IDRs)^2^. IDRs are unstructured, high-flexibility regions and are frequently enriched with specific amino acids (AAs), which may form weak but abundant interactions^3,4^. They include π-π stacking^5-7^, cation-π interactions^8-10^, electrostatic interactions^11-15^, and hydrophobic interactions^16-19^. These interactions not only drive scaffold proteins^2^ to form condensates but also mediate the specific partitioning of numerous client proteins^2,20-25^, which are not in a phase-separated state within the condensates because they lack the ability to undergo phase separation or fail to reach the necessary concentration threshold^26^.

Besides the simple partitioning or exclusion, researchers have started to tackle the factors affecting the coexistence of condensates, such as their mixing or demixing. A few mixing^27-34^ and demixing pairs^30,33-35^ have been identified, and various mechanisms that influence these behaviors have also been proposed, including interfacial tension^12,36-42^, the nature of forces driving condensate formation^43-45^, condensate material properties^46,47^, and the inherent competition between homotypic and heterotypic interactions^35,48-51^. Generally, weakened homotypic or enhanced heterotypic interactions promote mixing, and vice versa^48,49^. Additionally, we hypothesize that variations in the composition of phase separation-critical AAs among different IDRs lead them to preferentially form homotypic or heterotypic interactions, thereby determining whether their condensates mix or demix. This mechanism is not mutually exclusive of any of the aforementioned mechanisms.

In recognition that the previous studies were largely based on isolated cases, we carried out a systematic investigation to understand how AA composition influences the coexistence behavior of different condensates. We selected 28 IDRs that met two key criteria: (1) the ability to form condensates and (2) the formation of condensates driven by distinct forces, which are macroscopically evident through the inclusion of diverse AAs promoting phase separation, including serine^52^, aromatic residues^8,33,53-55^, and charged residues^13,29,56,57^. To ensure rapid and controllable formation of condensates by these IDRs, while preventing condensate aging, we employed the Cry2-based Optodroplet system^58^ and the iLID/SspB-based Corelets system^59^. We found that IDRs rich in serine or aromatic AAs tend to form miscible condensates. The presence of both serine and aromatic AAs generally enhances miscibility. In contrast, condensates formed by IDRs with high charge levels are typically immiscible, overriding the miscibility promoted by serine. Furthermore, phosphorylation of serine or tyrosine, which increases charge levels, plays a critical role in switching condensate miscibility. For example, condensates formed by the serine-rich SRSF1 were miscible with Pol II condensates until hyperphosphorylation increased its charge levels, impeding this miscibility. We also discovered that miscibility between transcription factors (TFs) and Pol II condensates crucially regulates transcription. With both artificial and modified native TFs, a high ratio of serine enhances miscibility and transactivation activity, whereas high levels of charge diminish these effects.

## Results

### The Selected IDRs Can Form Condensates

To explore the mechanisms underlying the miscibility of IDR condensates, we selected 28 representative IDRs (Figure S1A) known for their phase separation capabilities^7,10,14,52,54,60-70^. We analyzed 26 AA patterns in these IDRs (Figure S1B, Table S1): the ratios of the 20 standard AAs; the ratios of charged (EDKRH), negatively charged (ED), positively charged (KRH), and aromatic (FYW) AAs; the NCPR (net charge per residue); and the hydropathy values. We found that these IDRs are enriched with charged, aromatic, and some hydrophilic AAs but are deficient in hydrophobic AAs (C, I, L, V, A, and M).

We then analyzed the AA patterns in these IDRs using Spearman’s correlation. We observed a strong positive correlation between the ED and KRH ratios (Figure S1C) and hence between the combined EDKRH ratio and either ED or KRH ratios (Figure S1D-E). Moreover, we observed a strong positive correlation between the primary contributor to the aromatic content. Additionally, the FYW ratio negatively correlates with the EDKRH ratios (Figure S1G), providing an opportunity to independently investigate how electrostatic and π-based interactions influence the miscibility of condensates.

To enable homotypic and dynamic condensates for tractable interpretation, we used the two well-established light-control systems: the Cry2-based Optodroplet system^58^ and the iLID/SspB-based Corelets system^59^. Initially, we created 28 IDRs-mCherry-Cry2 constructs (Figure S2A), and all the IDRs except hnRNPK formed condensates upon blue light stimulation in cells (Figure S2B). Similarly, we generated 28 IDRs-GFP-iLID-T2A-SspB-FTH1 constructs (Figure S2C), and all except the BRD4 construct formed condensates upon blue light stimulation in cells (Figure S2D). Importantly, although hnRNPK in the Optodroplet system and BRD4 in the Corelets system did not form condensates, we could still assess their miscibility or immiscibility with other IDRs-formed condensates by using hnRNPK in the Corelets system and BRD4 in the Optodroplet system.

To assess the orthogonality between the Optodroplet and Corelets systems, and to eliminate any potential non-specific colocalization stemming from their backbones, we conducted the following experiments. We co-expressed the 28 IDRs-mCherry-Cry2 constructs with GFP-iLID-T2A-SspB-FTH1 and observed that GFP-iLID/SspB-FTH1 was not recruited into any IDRs-mCherry-Cry2 condensates (Figure S2E-F). Likewise, co-expression of the 28 IDRs-GFP-iLID-T2A-SspB-FTH1 constructs with mCherry-Cry2 showed no significant recruitment of mCherry-Cry2 into any IDRs-GFP-iLID/SspB-FTH1 condensates (Figure S2G-H). These findings confirm that the Optodroplet and Corelets systems are orthogonal to each other and to the selected IDRs, allowing for a comprehensive evaluation of pairwise condensate miscibility.

### Characterizing the Diversity of IDR Condensate Miscibility

Next, we delved deeper into the miscibility patterns among these condensates. We co-expressed two distinct constructs, each comprising a unique IDR linked to a different light-responsive backbone. Prior to capturing fluorescence images, the cells were exposed to blue light stimulation for 30-60 seconds, which was sufficient to allow IDR condensation, yet short enough to prevent potential hardening of the using the Pearson correlation coefficient, classifying those with a coefficient above 0.5 as miscible (Figure 1A) and those below 0.5 as immiscible (Figure 1C).

**Figure 1.**
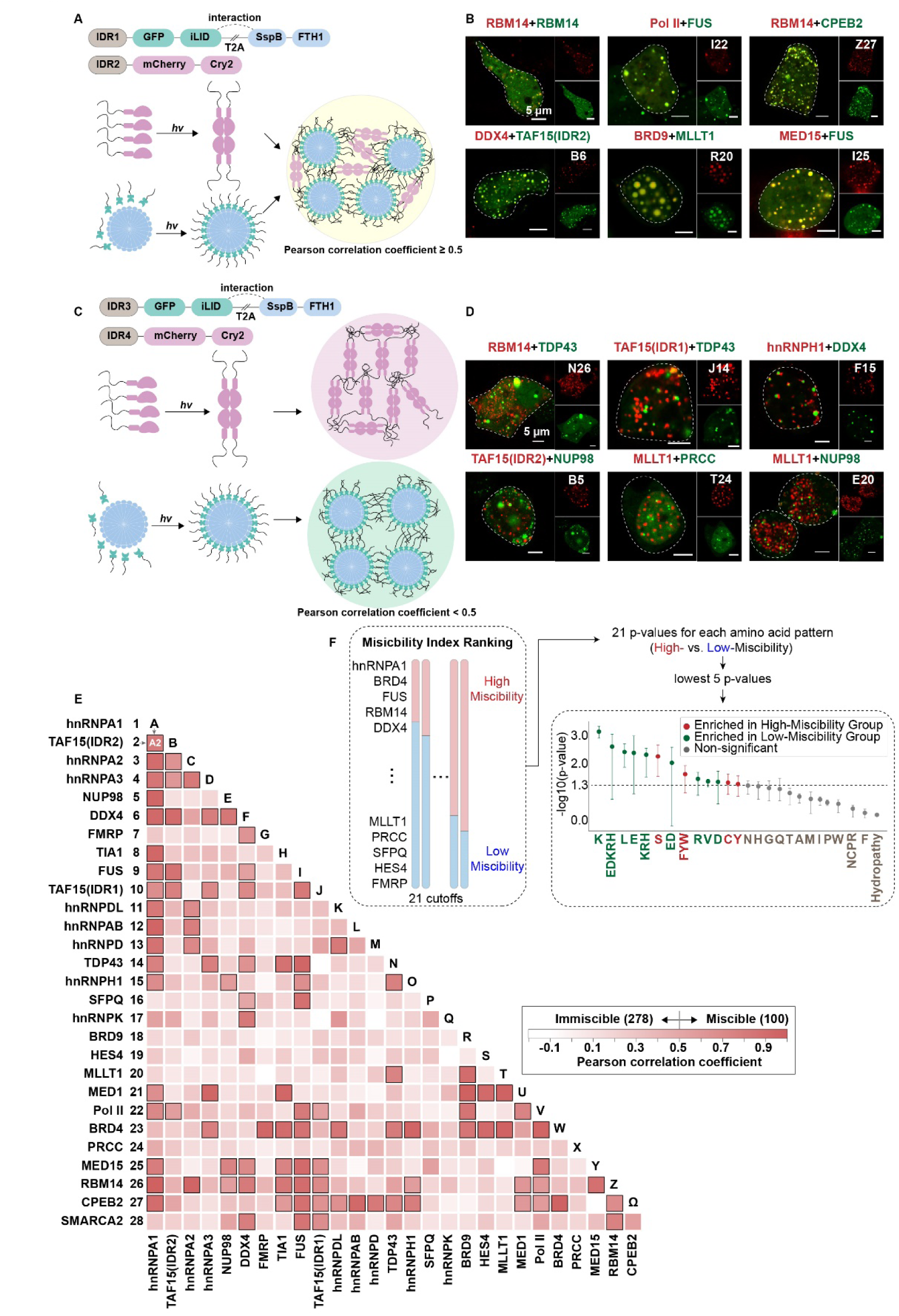
Analysis of IDR Condensate Formation. (A-B) Schematics and fluorescence images showing miscible condensates. The labels at the top-right corner of each image correspond to the row and column from Figure 1E. (C-D) Schematics and fluorescence images showing immiscible condensates. The labels at the top-right corner of each image correspond to the row and column from Figure 1E. (E) Heatmap of Pearson correlation coefficients to assess miscibility among the indicated pairs of IDR condensates. Squares outline in black signify miscible pairs with a Pearson correlation coefficient of 0.5 or higher. The matrix uses numbers 1-28 to label rows and letters A-Ω to label columns for easily finding the pairwise, such as A2 for the TAF15(IDR2)/hnRNPA1 pair, consistent with labels used in Figures S3. (F) The flow chart illustrating the process of analyzing the relationship between amino acid patterns and the miscibility of IDR condensates. It begins by categorizing the 28 IDR into high-miscibility and low-miscibility groups based on their Miscibility Index, with cutoffs varying from 4 to 24. The analysis encompasses 26 amino acid patterns, assessing their occurrences across these groups at different cutoffs, and generating 21 p-values for each pattern. The significance of each pattern is then established by selecting the lowest five p-values.

We first explored condensates formed by identical IDRs with different light-responsive backbones. All 28 IDR condensate pairs are miscible, as exemplified by the RBM14/RBM14 condensates (Figure 1B). We then extended our study to examine the miscibility of heterotypic IDR pairings, and identified 100 miscible pairs (Figures 1E and S3, Table S2). Five representative examples are shown, including the positive control, Pol II/FUS^33^, and four newly identified cases (Figure 1B). Simultaneously, we identified 278 immiscible pairs (Figure 1E, Table S2). Six representative examples are illustrated in Figure 1D, including two previously reported cases^30^, RBM14/TDP43 and TAF15(IDR1)/TDP43, and four newly identified ones.

The availability of the comprehensive set of images and sequences (Figure 1E, Tables S1 and S2) allowed us to interrogate how the AA composition influences the miscibility of IDR condensates. We first calculated a "Miscibility Index" for each IDR by averaging its Pearson correlation coefficients with 27 others (Table S2). We ranked these IDRs based on their Miscibility Index and split them into a high-miscibility group and a low-miscibility group by setting cutoffs, for instance, the top 4 IDRs are placed in the high-miscibility group, and the rest are in the low-miscibility group. This process was repeated with different cutoffs, up to 24, creating 21 different ways to categorize the IDRs (Figure 1F). Next, we used the 26 AA patterns in Figure S1B (the 20 standard AAs, FYW, EDKRH, KRH, ED, hydropathy, and NCPR) to assess statistically significant differences between the high- and low-miscibility groups. Each pattern was evaluated across 21 cutoffs, generating 21 p-values. The significance for each pattern was determined by the lowest five p-values. Our findings reveal that S, Y, and FYW are significantly enriched in the high-miscibility group. Conversely, nearly all charge-related AAs (E, D, K, R, ED, KRH, and EDKRH) are significantly enriched in the low-miscibility groups of IDR condensates (Figure 1F).

In summary, of the 378 IDR condensate pairs, 278 exhibited immiscible tendencies, while 100 showed miscible behaviors. Additionally, serine and aromatic AAs appear to enhance condensate miscibility, whereas charged AAs may play a negative role. Next, we will explore the effects of these composition features on the miscibility of condensates.

### Serine-rich Condensates Tend to Be Miscible

First, we explored the effect of serine residues on the miscibility of condensates. We ranked the 28 IDRs based on their S ratio, categorizing the top 50% as S(Rich) and the bottom 50% as S(Poor) (Figure 2A), between which only the serine ratio was significantly higher in the S(Rich) group (Figure 2B). Further analysis indicated that the miscibility of S(Rich) condensates was significantly higher than that of S(Poor) condensates (Figure 2C-E). This observation aligns with the notion that serine can weaken homotypic interactions^71,72^ by enhancing effective solvation volume^6^, thereby lowering the barrier for condensate mixing^51^.

**Figure 2.**
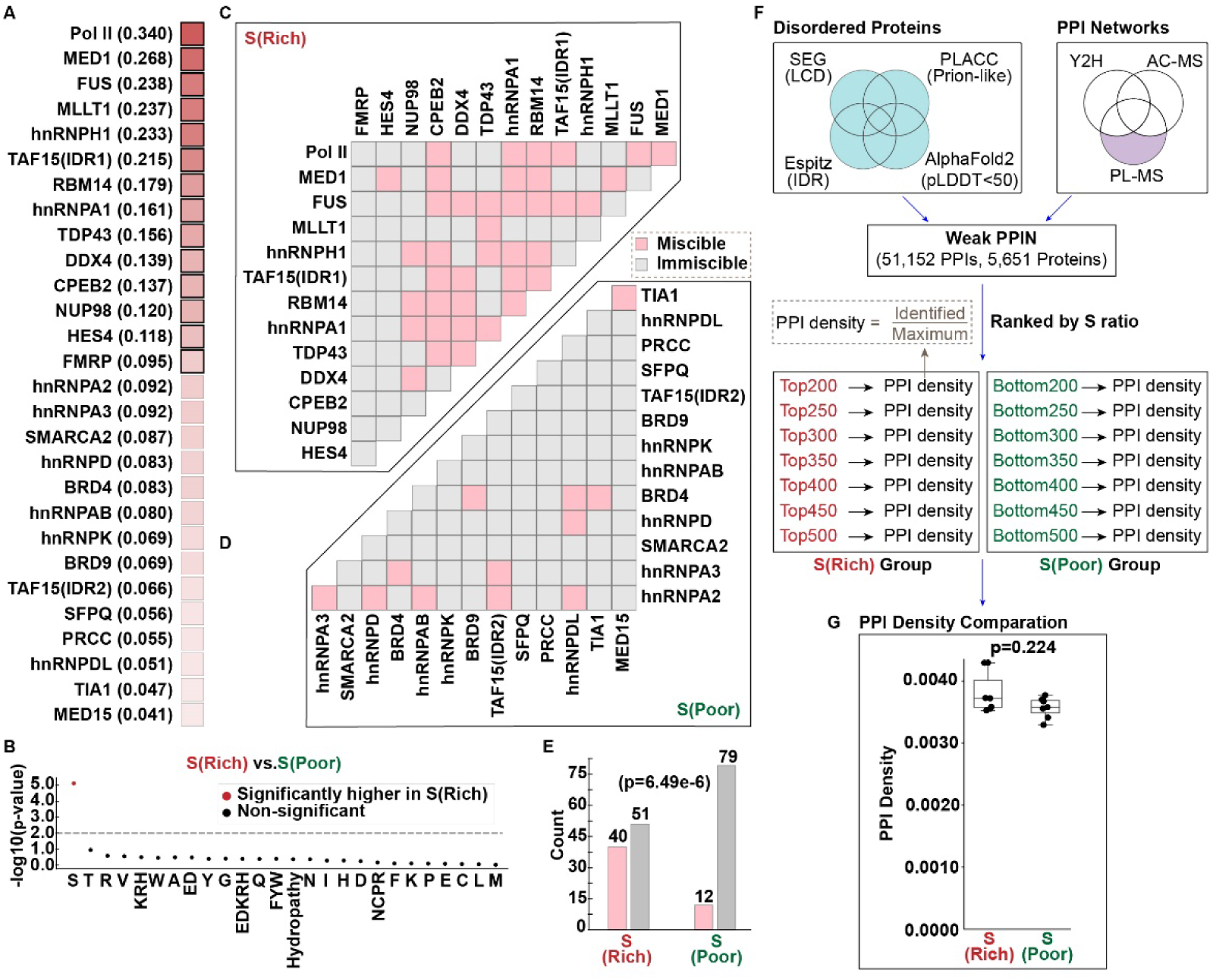
Serine-rich IDR Condensates Show High Miscibility. (A) Heatmap displaying the ranking of 28 IDRs based on their serine (S) ratios. The top 50% of IDRs are classified as S(Rich). The remaining IDRs are categorized as S(Poor). The S ratios are in brackets at the right of each IDR name. (B) Plot presenting statistical differences in the 26 AA patterns between IDRs in the S(Rich) group and IDRs in the S(Poor) group. (C-E) Matrices depicting miscible (soft pink) and immiscible (light gray) pairs of condensates formed by S(Rich) IDRs and S(Poor) IDRs, followed by a quantification of the data, analyzed using Fisher’s exact test. (F) The flowchart details the process used to analyze differences in PPI densities between the S(Rich) group and the S(Poor) group within weak protein-protein interaction network (PPIN). Disordered regions include low-complexity domains (LCDs) as predicted by SEG, prion-like domains identified by PLAAC, intrinsically disordered regions (IDRs) by Espritz, and regions with a pLDDT score below 50 in AlphaFold2. The analysis then focuses on PPIs that were detected using Proximity-Labeling Mass Spectrometry (PL-MS) but not by Yeast Two-Hybrid (Y2H) or Affinity Capture-MS (AC-MS). These PPIs, identified among these disordered proteins, form the basis of weak PPIN. PPI densities were calculated by comparing the identified number of PPIs against the theoretical maximum. These proteins were ranked according to their S ratio. Groups containing the top 200, 250, 300, 350, 400, 450, and 500 proteins were classified as S(Rich) groups, while those with the bottom 200, 250, 300, 350, 400, 450, and 500 were classified as S(Poor) groups. (G) Comparison of PPI densities between S(Rich) proteins and S(Poor) proteins. Mann-Whitney U test was performed.

Beyond weakening homotypic interactions, we explored whether serine might also promote miscibility by enhancing heterotypic interactions. To explore this, we focused on weak Protein-Protein Interaction Network (PPIN). We first identified 5,651 human proteins that contain at least one IDR of 50 or more AAs, which are also recorded in Proximity-labeling Mass Spectrometry (PL-MS) databases but absent from Yeast Two-Hybrid (Y2H) and Affinity Capture-MS (AC-MS) databases (Table S3). The PPIs among these proteins are considered weak. We ranked these proteins according to their serine ratios in IDRs and selected the top 200 proteins. Then we calculated their PPI densities by comparing the identified number of PPIs against the theoretical maximum. In total, 14 cutoffs—top and bottom 200, 250, 300, 350, 400, 450, and 500—were established to calculate the PPI densities, yielding seven densities for each of the top (S(Rich)) and bottom (S(Poor)) groups (Figure 2F, Table S3). The results revealed no significant differences in PPI densities between the S(Rich) and S(Poor) groups (Figure 2G), indicating that serine-rich proteins did not display a tendency to increase heterotypic interactions under the analyzed conditions.

These data suggest that serine-rich condensates are more miscible, likely because serine tends to weaken homotypic interactions rather than to enhance heterotypic ones.

### Aromatic-rich Condensates Tend to Be Miscible

We then examined the impact of aromatic AAs (FYW) on the miscibility properties of condensates. We ranked the 28 IDRs based on their FYW ratio, categorizing the top 50% as FYW(Rich) and the bottom 50% as FYW(Poor) (x-axis in Figure S4A). Our findings showed that the FYW(Rich) group had a significantly higher miscibility than the FYW(Poor) group (Figure 3A-C), suggesting that aromatic AAs may enhance condensate miscibility. However, our analysis of AA patterns revealed that, beyond FYW and Y, other AA patterns also displayed significant differences between the FYW(Rich) and FYW(Poor) groups (Figure S4B). Therefore, we refined our IDRs by removing those IDRs rich in EDKRH from the FYW(Rich) group and excluding some IDRs from the FYW(Poor) group due to having extremely low or zero ratios of L, V, I, and A (Figure S4A-B). After this adjustment, only the aromatic AAs ratio remained significantly different between the newly defined groups (Figure S4C). Additionally, the miscibility of condensates in the FYW(Rich-filtered) group remained significantly higher than that in the FYW(Poor-filtered) group (Figure S4D-F). The analysis of PPI densities in weak PPIN demonstrated that proteins rich in aromatic AAs exhibited significantly higher interaction densities among themselves compared to those lacking these AAs (Figure 3D, Table S3). These data suggest that a high ratio of aromatic AAs promotes the mixing of condensates, likely through the formation of heterotypic π-based interactions.

**Figure 3.**
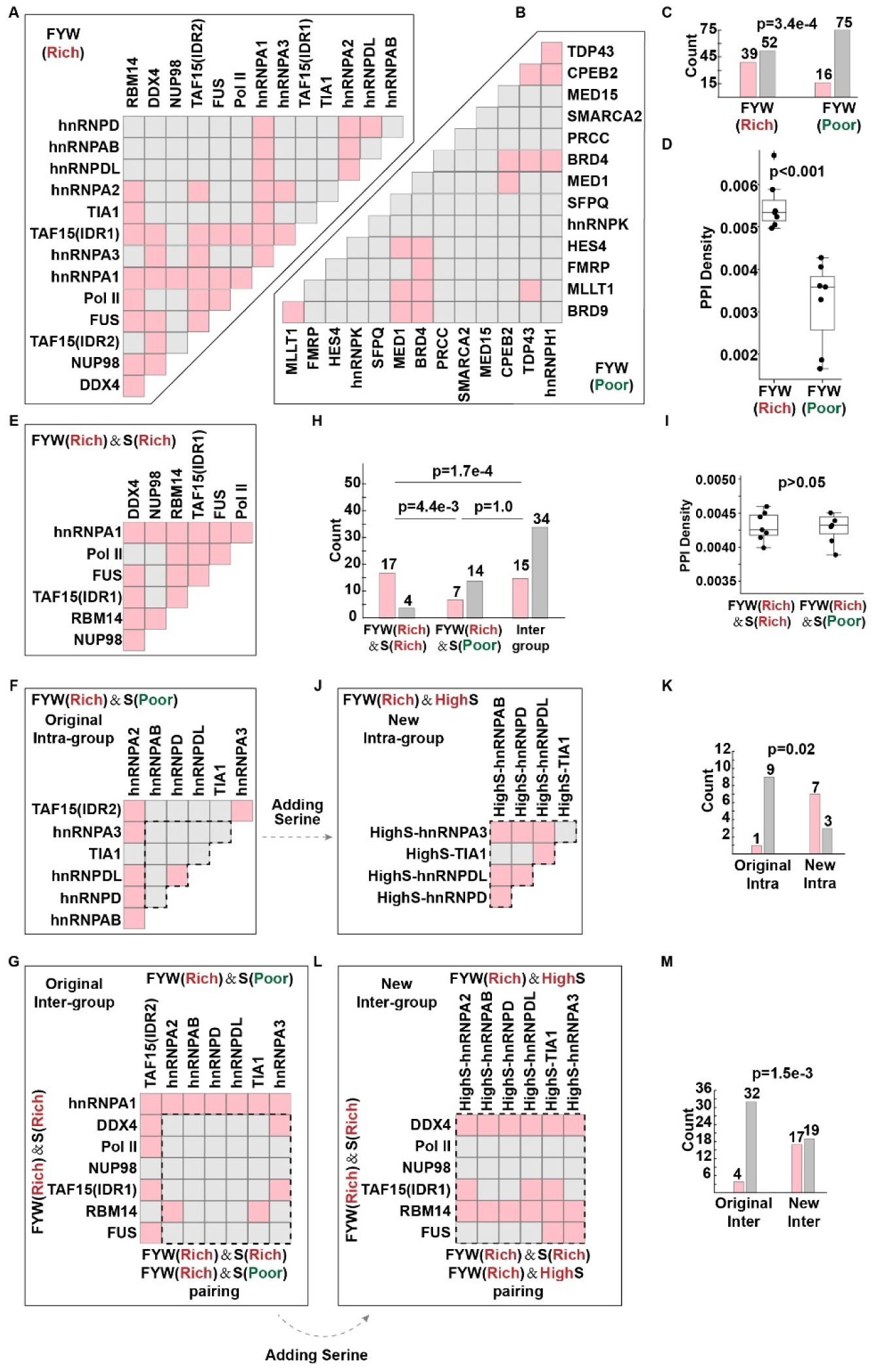
Aromatic-rich Condensates Tend to Be Miscible. (A-B) Matrices showing miscible (soft pink) and immiscible (light gray) pairs of condensates: (A) formed by FYW(Rich) IDRs and (B) formed by FYW(Poor) IDRs. (C) Quantification of the data from Panels A and B. Fisher’s exact test was performed. (D) Comparison of PPI densities between FYW(Rich) proteins with FYW(Poor) proteins. Mann-Whitney U test was performed. (E-F) Matrices showing miscible (soft pink) and immiscible (light gray) pairs of condensates: (E) formed by FYW(Rich)&S(Rich) IDRs and (F) formed by FYW(Rich)&S(Poor) IDRs. Dashed line highlights the IDRs selected for fusion with the serine-rich peptide, as detailed in Panel J. (G) Matrix displaying miscible (soft pink) and immiscible (light gray) condensates formed by pairing FYW(Rich)&S(Rich) IDRs with FYW(Rich)&S(Poor) IDRs. Dashed line highlights the IDRs selected for fusion with the serine-rich peptide, as detailed in Panel L. (H) Quantification of the data from Panels E-G. Fisher’s exact test was performed. (I) Comparison of PPI densities between FYW(Rich)&S(Rich) proteins with FYW(Rich)&S(Poor) proteins. Mann-Whitney U test was performed. (J) Matrix showing miscible (soft pink) and immiscible (light gray) pairs of condensates formed by FYW(Rich)&HighS IDRs. (K) Quantification of the data from Panels F and J. Fisher’s exact test was performed. (L) Matrix displaying miscible (soft pink) and immiscible (light gray) condensates formed by pairing FYW(Rich)&S(Rich) IDRs with FYW(Rich)&HighS IDRs. (M) Quantification of the data from Panels G and L. Fisher’s exact test was performed.

It is notable that nearly 60% (52 out of 91) of the pairs in the FYW(Rich) group still exhibited limited miscibility (Figure 3A), which may be due to strong homotypic interactions. Given that serine can reduce homotypic interactions^6^, we hypothesized that a high serine ratio might enhance the miscibility of aromatic-rich condensates. We then re-ranked the FYW-rich IDRs based on their serine ratio, categorizing the top 50% as FYW(Rich)&S(Rich) and the bottom 50% as FYW(Rich)&S(Poor) (Figure S4G). Further analysis showed that only serine ratio was significantly higher in the FYW(Rich)&S(Rich) group (Figure S4H). Notably, this group also displayed a significantly higher miscibility compared to both the FYW(Rich)&S(Poor) group and the intergroup (Figure 3E-H). Furthermore, weak PPIN analysis revealed no significant difference in PPI densities between the FYW(Rich)&S(Rich) and FYW(Rich)&S(Poor) groups (Figure 3I, Table S3), implying that serine did not facilitate these aromatic-rich proteins to form more heterotypic interactions.

To further explore the role of serine in enhancing the miscibility of aromatic-rich condensates, we engineered a serine-rich disordered peptide named HighS (Figure S4I). The HighS peptide fused to mCherry and Cry2 neither formed condensates nor partitioned into condensates formed by FYW(Rich)&S(Rich) IDRs (Figure S4J). We then fused HighS with the FYW(Rich)&S(Poor) IDRs, forming a new group called FYW(Rich)&HighS, which was enriched in FYW and serine. The miscibility of condensates in this new group was significantly higher compared to the original FYW(Rich)&S(Poor) group (Figures 3F, 3J-K, S4K). Moreover, miscibility was significantly higher in the new intergroup (FYW(Rich)&S(Rich) and FYW(Rich)&HighS) compared to the original (Figure 3G, 3L-M, S4L).

In conclusion, condensates rich in aromatic amino acids exhibit high miscibility, and increasing the proportion of serine further enhances this miscibility, not by facilitating heterotypic interactions, but likely by reducing homotypic interactions.

### Charge-rich Condensates Tend to Be Immiscible

We then investigated how charge properties affect the miscibility of condensates. Initially, we focused on IDRs with positive NCPR and those with negative NCPR, which theoretically could engage in heterotypic electrostatic interactions. After ranking the 28 IDRs by their NCPR, we selected the six highest positive values and the four lowest negative values (Figure 4A). Contrary to expectations, no miscible pairs were found, regardless of whether positive IDRs were paired with negative ones or IDRs were grouped by the same charge type (Figure 4B-D). Similarly surprising, weak PPIN analysis revealed that there was no significant difference in PPI densities between the two groups: (positive paired with negative) versus (positive paired with neutral) (Figure 4E, Table S3), suggesting that proteins with opposite NCPRs do not have a general tendency to form heterotypic interactions, potentially explaining why condensates composed of IDRs with opposing charges tend to resist mixing.

**Figure 4.**
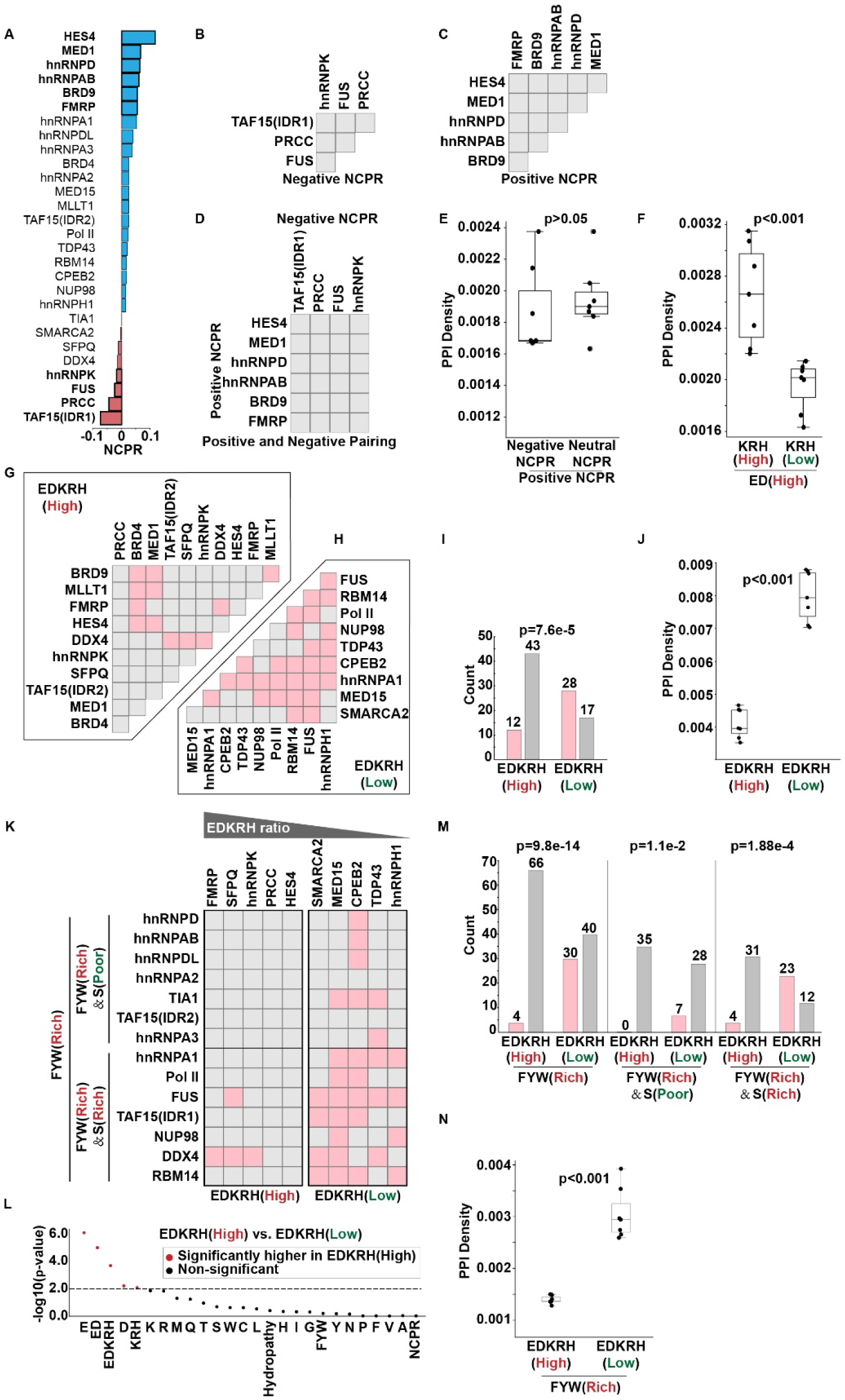
Charge-rich Condensates Show High Immiscibility. (A) Bar chart displaying the NCPR values for the 28 IDRs, ranked from highest to lowest. The 10 IDRs in bold text were selected for further analysis. (B-D) Matrices showing miscibility (soft pink) and immiscibility (light gray) of condensates: (B) formed by pairs of IDRs with negative NCPR, and (C) formed by pairs of IDRs with positive NCPR, (D) formed by IDRs with positive NCPR paired with those with negative NCPR. Only immiscible cases were observed in all matrices. (E) Comparison of PPI densities in protein pairings based on NCPR. Mann-Whitney U test was performed. (F) Comparison of PPI densities between paired proteins categorized by charged AA ratios: ED(High) paired with KRH(High) vs. ED(High) paired with KRH(Low). Mann-Whitney U test was performed. (G-H) Matrices showing miscibility (soft pink) and immiscibility (light gray) of condensates: (G) formed by pairs of IDRs with high EDKRH ratio and (H) formed by pairs of IDRs with low EDKRH ratio. (I) Quantification of the data from Panels G and H. Fisher’s exact test was performed. (J) Comparison of PPI densities between EDKRH(High) proteins with EDKRH(Low) proteins. Mann-Whitney U test was performed. (K) Matrix showing miscibility (soft pink) and immiscibility (light gray) of condensates formed by FYW(Rich), FYW(Rich)&S(Rich) or FYW(Rich)&S(Poor) IDRs paired with those formed by EDKRH(High) or EDKRH(Low) IDRs. (L) Plot showing statistical differences in the 26 AA patterns between IDRs EDKRH(High) IDRs and EDKRH(Low) IDRs. (M) Quantification of the data from Panel K. Fisher’s exact test was performed. (N) Comparison of PPI densities in protein pairings involving FYW(Rich) proteins: paired with EDKRH(High) and EDKRH(Low). Mann-Whitney U test was performed.

Beyond NCPR, we assessed the impact of absolute ratios of positively charged (KRH) and negatively charged (ED) amino acids within IDRs on the formation of heterotypic interactions. Weak PPIN analysis indicated that ED(High) proteins form significantly more heterotypic interactions with KRH(High) than with KRH(Low) (Figure 4F, Table S3). However, due to a strong positive correlation between the ED and KRH ratios among the IDR set (Figure S1C), there were too few IDRs containing only high ED or KRH content, preventing us from evaluating whether these heterotypic interactions facilitated effective condensate mixing.

As a result, we shifted our focus to IDRs rich in both ED and KRH, which are theoretically capable of forming both homotypic and heterotypic electrostatic interactions. To minimize the influence from aromatic AAs, we excluded IDRs with a high FYW ratio (top 25%) (Figure S5A). We then ranked the remaining 21 IDRs by their EDKRH ratios, dividing them into two nearly equal groups: EDKRH(High) and EDKRH(Low) (Figure S5A). Analysis of the 26 AA patterns revealed that only the ratios of charged AAs were significantly different between these groups (Figure S5B). Moreover, the miscibility of condensates in the EDKRH(High) group was significantly lower than in the EDKRH(Low) group (Figure 4G-I). Similarly, weak PPIN analysis revealed that PPI densities were significantly lower in the EDKRH(High) group compared to the EDKRH(Low) group (Figure 4J, Table S3). These findings suggest that condensates rich in charged AAs tend to disfavor mixing, likely due to their limited ability to form heterotypic interactions.

Previous studies have emphasized the importance of charged AA distribution patterns in affecting the homotypic or heterotypic interaction among IDRs^73-81^. We then investigated whether these patterns also influence condensate miscibility. Upon analyzing our selected IDRs, we identified four as ‘charge-blocky’ and seven as ‘charge-dispersed’ (Figure S5C). Further analysis showed that the number of charge blocks was the only significantly different feature between these two groups (Figure S5D). The assessment of miscibility showed that condensates formed by charge-blocky IDRs tended to mix with each other (Figure S5E), while those formed by charge-dispersed IDRs resisted mixing both among themselves (Figure S5F) and with those formed by charge-blocky IDRs (Figure S5G-H).

We also assessed whether charge-rich condensates still prefer demixing from other types of condensates, such as aromatic-rich condensates. We ranked the 28 IDRs by their FYW and EDKRH ratios, categorizing the top 50% in FYW ratio as FYW(Rich) and the bottom 50% as FYW(Poor) (Figure S4A). FYW(Rich) IDRs were further subdivided based on serine content: FYW(Rich)&S(Rich) and FYW(Rich)&S(Poor) (Figure S4G), while FYW(Poor) IDRs were classified into EDKRH(High) and EDKRH(Low) groups (Figure 4K). Analysis of the 26 AA patterns revealed that the ratio of EDKRH was significantly different between the EDKRH(High) and EDKRH(Low) groups (Figure 4L). FYW(Rich) condensates, irrespective of serine content, generally resisted mixing with charge-rich condensates (EDKRH(High)) (Figure 4M). Furthermore, weak PPIN analysis revealed that FYW(Rich) proteins formed fewer PPIs with EDKRH(High) proteins compared to EDKRH(Low) (Figure 4N, Table S3). These findings indicate that charge-rich condensates resist mixing with aromatic-rich condensates, probably because of their restricted capacity to engage in heterotypic interactions.

In conclusion, condensates formed by highly charged IDRs tend to be highly immiscible.

### Charge Overwrites Serine-enhanced Miscibility

We next investigated whether a high ratio of serine could enhance the miscibility of condensates formed by charge-rich IDRs. We initially ranked the 28 IDRs based on their serine and EDKRH ratios, dividing them into four groups: EDKRH(Low)&S(Poor), EDKRH(Low)&S(Rich), EDKRH(High)&S(Poor), and EDKRH(High)&S(Rich) (Figure S5I). A comparative analysis of the amino acid patterns between the EDKRH(High)&S(Poor) and EDKRH(High)&S(Rich) groups revealed that the serine content was the only significant variable (Figure S5J). However, our assessment of miscibility between these groups showed no significant differences (Figure 5A, 5B, 5D). We then fused the serine-rich peptide with the EDKRH(High)&S(Poor) IDRs to assess if increased serine content would enhance miscibility. The resulting condensates, however, still showed a tendency to resist mixing (Figures 5C, S5K). Furthermore, weak PPIN analysis revealed that PPI densities were significantly lower in the EDKRH(High)&S(Rich) group compared to the EDKRH(High)&S(Poor) group (Figure 5E, Table S3), suggesting that serine does not facilitate, and may even hinder, the formation of heterotypic interactions among charge-rich proteins. Overall, our findings indicate that a high serine ratio does not improve the miscibility of condensates formed by charge-rich IDRs.

**Figure 5.**
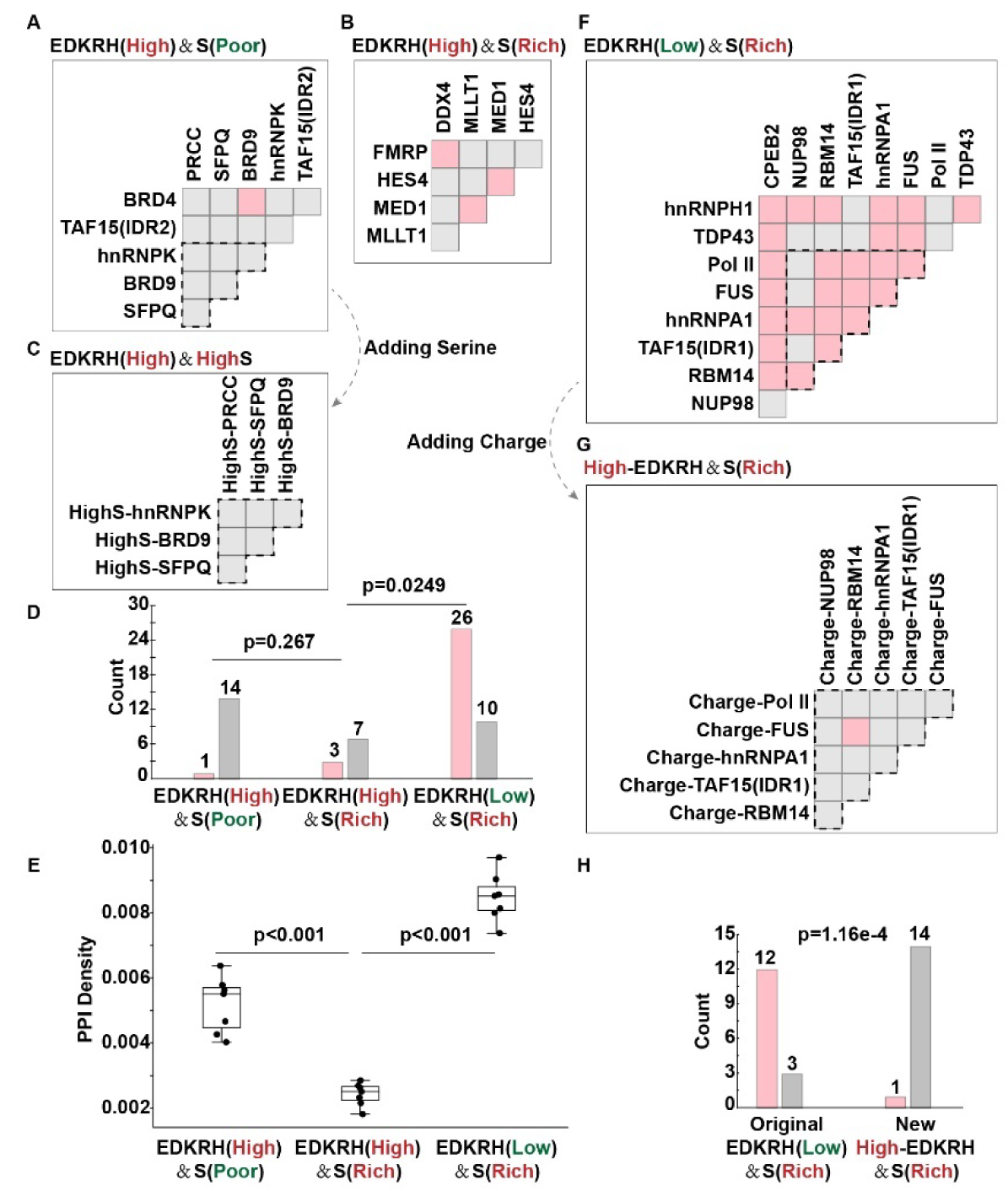
Charges Are Dominant Contributors to Immiscibility. (A-C) Matrices showing miscible (soft pink) and immiscible (light gray) pairs of condensates: (A) formed by EDKRH(High)&S(Poor) IDRs. Dashed line highlights the IDRs selected for fusion with the serine-rich peptide, as detailed in Panel C, (B) formed by EDKRH(High)&S(Rich) IDRs, and (C) formed by EDKRH(High)&HighS IDRs. (D) Quantification of the data from Panels A, B, and F. Fisher’s exact test was performed. (E) Comparison of PPI densities among proteins: EDKRH(High)&S(Poor) proteins vs. EDKRH(High)&S(Rich) proteins vs. EDKRH(Low)&S(Rich) proteins. Mann-Whitney U test was performed. (F-G) Matrices showing miscible (soft pink) and immiscible (light gray) pairs of condensates: (F) formed by EDKRH(Low)&S(Rich) IDRs. Dashed line highlights the IDRs selected for fusion with the charge-rich peptide, as detailed in Panel G, and (G) formed by High-EDKRH&S(Rich) IDRs. (H) Quantification of the data from Panels F and G. Fisher’s exact test was performed.

Next, we investigated whether a high ratio of charged AAs can compromise the miscibility enhanced by a high serine ratio. We found that the condensates in the EDKRH(High)&S(Rich) group exhibited significantly lower miscibility compared to those in the EDKRH(Low)&S(Rich) group (Figure 5B, 5D, 5F). Analysis of the AA patterns revealed that the EDKRH ratio was the most significantly different feature between the two groups (Figure S5L). Moreover, results from weak PPIN analysis indicated that PPI densities were significantly lower among the proteins in the EDKRH(High)&S(Rich) group compared to the EDKRH(Low)&S(Rich) group (Figure 5E, Table S3), suggesting that proteins rich in serine and charges resist forming heterotypic interactions. To further examine the influence of charges in counteracting the miscibility enhancement by serine, we engineered a peptide, named High-EDKRH, rich in dispersed charged AAs that lacked the capacity to form condensates (Figure S5M-N). We found that the condensates formed by these new High-EDKRH&S(Rich) IDRs showed a reduced preference for mixing with each other (Figures 5G-H, S5O). These results suggest that high levels of charge can indeed compromise the miscibility enhanced by a high serine ratio.

In conclusion, high levels of charge are dominant contributors to the immiscibility of IDR condensates, overriding the positive contribution of serine-richness in promoting condensate miscibility.

### Phosphorylation Alters Protein Condensate Miscibility

We elucidated the roles of serine and aromatic amino acids, particularly tyrosine, in enhancing condensate miscibility, and identified the significant contribution of high charge levels to immiscibility. Notably, both serine and tyrosine can be phosphorylated, significantly increasing charge levels. We propose that phosphorylation is a critical regulator of condensate miscibility.

To investigate the impact of phosphorylation on protein phase separation, we analyzed 12,111 human disordered proteins, identifying 8,672 proteins that contain at least one residue phosphorylated at serine or tyrosine (Figure 6A, Table S4). Using SaPS, a phase-separation predictor, we assessed the phase separation potential of these proteins in both their native and phosphorylated states. Setting a SaPS score threshold of 0.5 to indicate the capability for phase separation, we found that proteins capable of phase separation exhibited significantly higher levels of phosphorylation as well (Figure 6B, Table S4). Additionally, phosphorylation notably increased the overall SaPS scores of these proteins (Figure 6C). Subsequent analysis revealed a strong positive correlation between the levels of protein phosphorylation and their propensity to undergo phase separation (Figure 6D, Table S4). These findings indicate that phosphorylation significantly influences protein phase separation propensity, potentially serving as the regulating switch in condensate miscibility.

**Figure 6.**
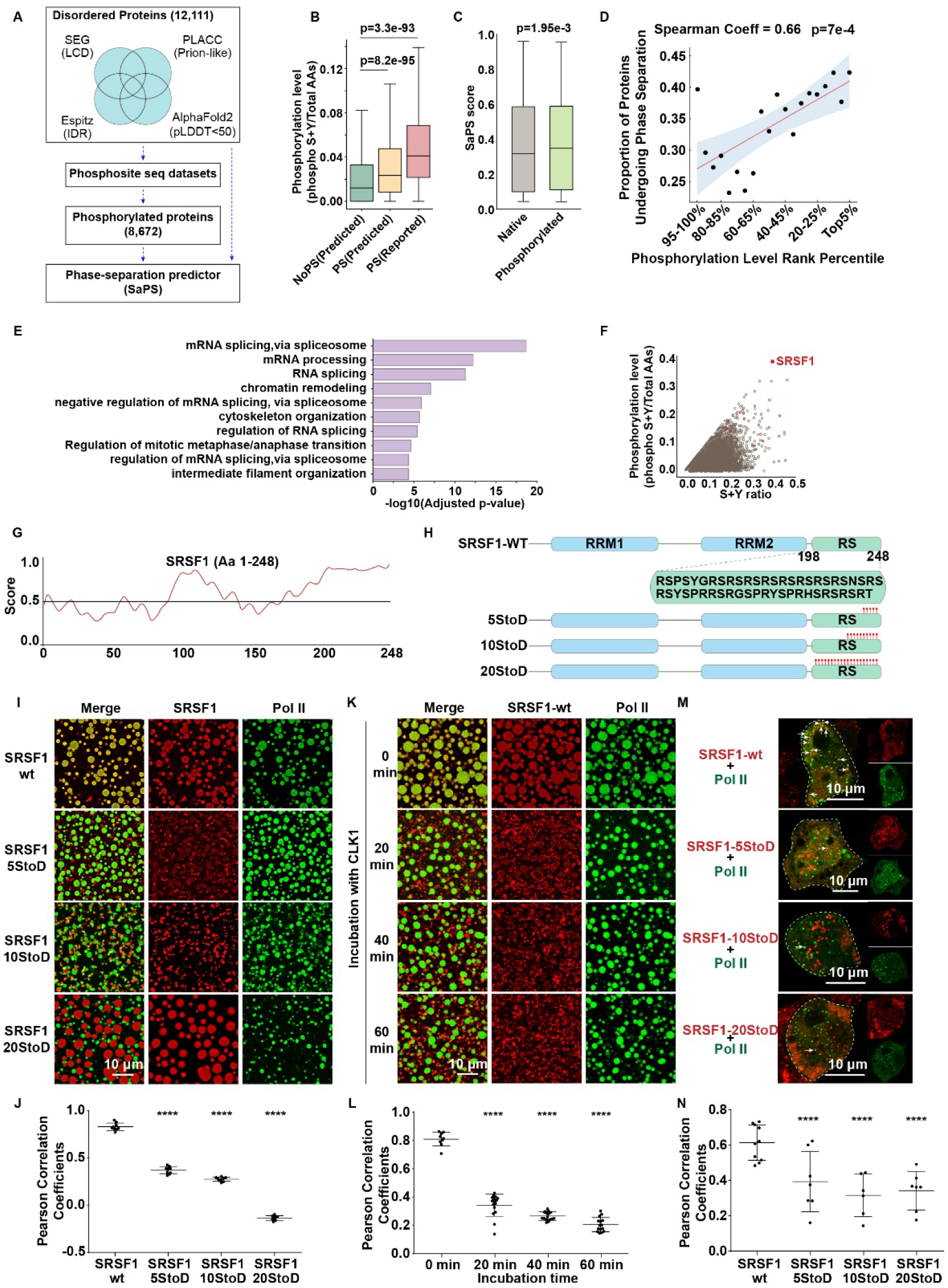
Phosphorylation Increases SRSF1 Charge and Reduces Miscibility with Pol II Condensates. (A) Identification of 12,111 human proteins with disordered regions (length ≥ 50 amino acids) using SEG, PLAAC, Espritz, and AlphaFold2. Of these, 8,672 contain at least one phosphorylated serine or tyrosine residue. SaPS was used to assess the phase separation potential of these proteins in both their native and phosphorylated states. (B) Comparison of phosphorylation levels at serine and tyrosine (S+Y) across proteins categorized by their potential to undergo phase separation. PS(Reported) includes proteins confirmed to undergo LLPS in databases like PhaSepDB 2.1, LLPSDB, and PhaSePro. PS(Predicted) covers proteins predicted to undergo phase separation with a SaPS score ≥0.5, while NoPS(Predicted) includes those with a SaPS score < 0.5. (C) Comparison of SaPS scores between proteins in their native and phosphorylated states. (D) Plot showing the correlation between phosphorylation levels and the proportion of proteins undergoing phase separation. Proteins are categorized on the x-axis by their phosphorylation level ranks, from the highest (top 5%) to the lowest (95-100%). The y-axis shows the ratio of proteins capable of phase separation (with a score ≥ 0.5). (E) Gene Ontology (GO) enrichment analysis was conducted on proteins with SaPS scores ≥0.5 and phosphorylation levels in the top 20%. A total of 822 proteins, containing 1,064 IDRs, were selected. (F) Scatter plot illustrating the relationship between the S+Y ratio and S+Y phosphorylation ratio across 5,322 IDRs from the 2,945 proteins with a SaPS score ≥ 0.5. The S+Y phosphorylation ratio is calculated as the number of phosphorylated S and Y residues relative to the total number of amino acids in the IDRs. Red circles highlight core mRNA splicing proteins, with SRSF1 exhibiting the highest S+Y phosphorylation ratio. (G) A disorder prediction for full-length SRSF1 by IUPred2A (https://iupred2a.elte.hu/). (H) To mimic different levels of phosphorylation, variants SRSF1-5StoD, SRSF1-10StoD, and SRSF1-20StoD were created with five, ten, and twenty serine or tyrosine residues in the RS domain replaced with negatively-charged AAs, indicated by red bars. (I-J) Fluorescence imaging (I) and quantitative co-localization analysis (J) of Pol II condensates with condensates formed by SRSF1-wt and various phosphomimetic SRSF1 variants in vitro. All proteins concentrations were 10 μM. (K-L) Time-course fluorescence imaging (K) and quantitative co-localization analysis (L) of Pol II condensates and condensates formed by wild-type SRSF1 after incubation with CLK1. (M-N) Fluorescence imaging (M) and quantitative co-localization analysis (N) of Pol II condensates with condensates formed by SRSF1-wt and various phosphomimetic SRSF1 variants, fused with Cry2-mCherry, in cells. The arrows in the merged images in Panel M indicate the co-localized condensates.

To determine which cellular processes are most profoundly influenced by phosphorylation-induced phase separation changes, we conducted a Gene Ontology (GO) enrichment analysis. We identified 2,945 proteins, each with a SaPS score above 0.5 and at least one phosphorylation site, containing a total of 5,322 IDRs. For further GO analysis, we focused on IDRs (1,064 IDRs from 822 proteins, as detailed in Table S4) with phosphorylation levels in the top 20%. Our results indicate a significant involvement of these proteins in various biological processes, particularly mRNA splicing (Figure 6E). Additionally, when we plotted the serine plus tyrosine (S+Y) ratio of these 5,322 IDRs versus their phosphorylation levels, we observed that Serine/Arginine-Rich Splicing Factors (SRSFs), critical components of the spliceosome^82^, had very high S+Y ratios and were also highly phosphorylated. Notably, SRSF1 exhibited the highest phosphorylated S+Y ratio (Figure 6F).

Considering reports that SRSF1 condensates can mix with Pol II condensates^31^, we propose that SRSF1 phosphorylation dynamically regulates the miscibility of SRSF1 with Pol II condensates by altering unmodified serine and tyrosine content, and charge levels. To mimic phosphorylation, we mutated serine and tyrosine residues to aspartic acids (Figure 6G-H, Document S1), decreasing the ratio of unmodified serine and tyrosine and increasing the ratio of charged AAs (Figure S6A). These modifications enhanced SRSF1’s phase separation capabilities (Figure S6B), suggesting that phosphorylation promotes its homotypic interactions. We then examined the impact of phosphorylation on the miscibility of SRSF1 condensates with Pol II. Our results show that while purified wild-type SRSF1 displayed high miscibility with Pol II condensates, increased phosphorylation levels led to decreased miscibility (Figure 6I-J). Similar trends were observed when SRSF1 was phosphorylated by CLK1 (Figures 6K-L, S6C), a kinase specific to the SRSF1 RS domain^83^. Moreover, when co-expressing Pol II with wild-type SRSF1 and its variants, wild-type SRSF1 condensates significantly colocalized with Pol II condensates, while phosphorylation-mimicking variants showed reduced miscibility (Figure 6M-N).

In conclusion, our research shows that phosphorylation crucially influences the miscibility of protein condensates by altering the levels of unmodified serine and tyrosine, and overall charge.

### Highly Charged TFs Exhibit Low Miscibility with Pol II Condensates and Reduced Transactivation

Finally, we investigated whether cells regulate functions such as transcription by altering the miscibility of condensates. To explore this, we engineered 27 artificial TFs, termed IDRs-SOX2. Each TF was created by fusing the DNA-binding domain of SOX2 (SOX2(DBD)) to one of the 27 IDRs, excluding the Pol II IDR. These constructs were individually co-transfected into 293T cells with a firefly luciferase reporter vector containing 26 tandem SOX2 binding motifs (Figure 7A). The dual-luciferase assays revealed variable effects on target gene expression: some IDRs-SOX2 fusions significantly enhanced expression, others had minimal impact, and a few even reduced it (Figure 7B, Table S5). Overall, miscibility between Pol II and IDR condensates, as measured by Pearson’s coefficient, demonstrated a positive correlation with the transactivation activity of the artificial IDRs-SOX2 TFs (Figure 7C).

**Figure 7.**
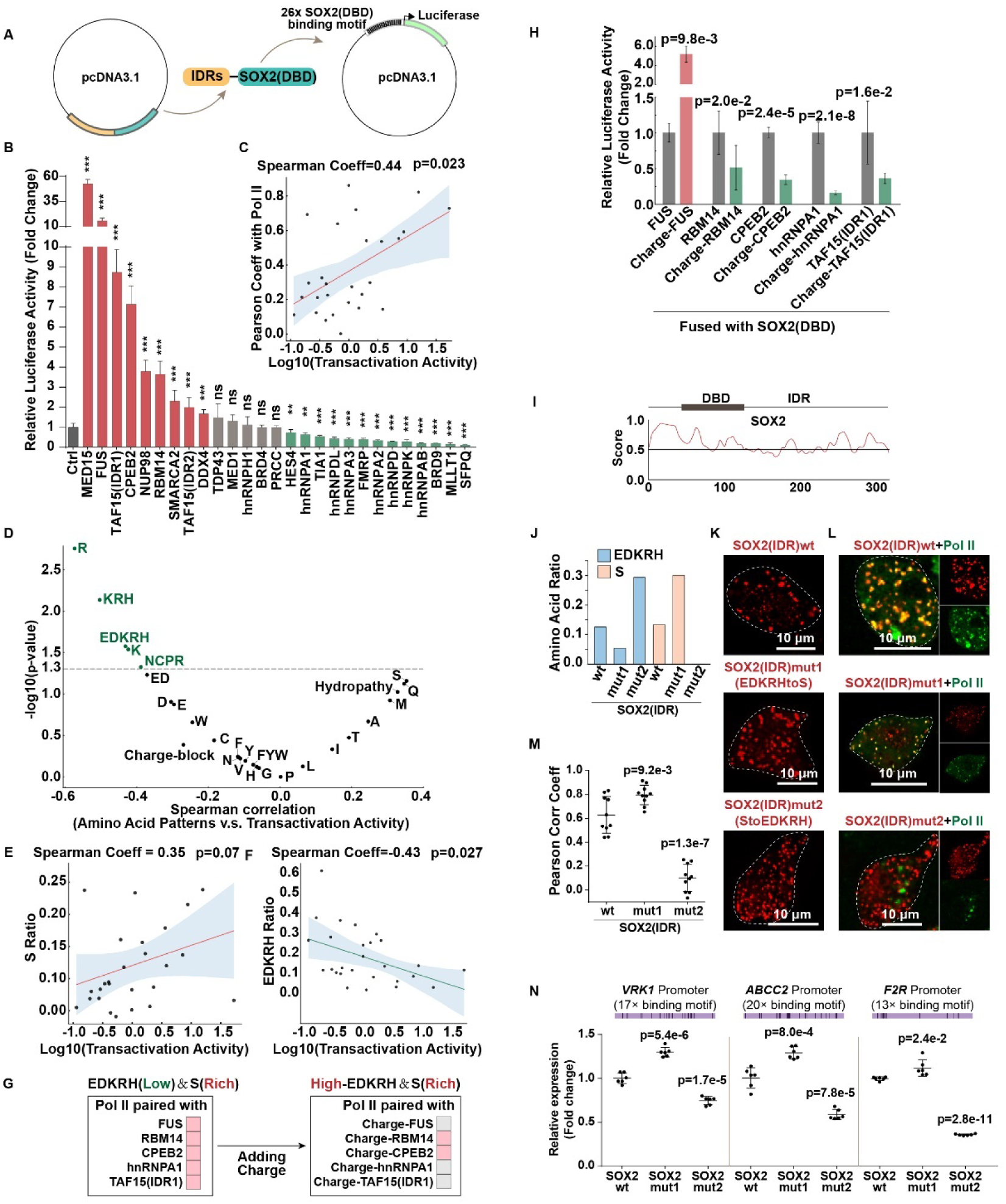
Highly Charged TFs Condensates Show Low Miscibility with Pol II Condensates and Decreased Transactivation. (A) Schematic diagram of the dual-luciferase assay. (B) Outcomes from dual-luciferase assays assessing the transcriptional activity of IDRs-SOX2 proteins. Bars colored red represent significant enhancement, whereas those colored green denote significant repression. (C) Spearman correlation analysis of the log10-transformed transactivation activity of 27 IDRs-SOX2 proteins versus their miscibility with Pol II condensates, indicated by Pearson coefficients. The red line shows the regression best-fit, and the shading indicates the 95% confidence interval. (D) Volcano plot presenting Spearman correlation analysis of amino acid patterns in 27 IDRs versus the transactivation activity of 27 IDRs-SOX2. Green points above the dashed line (p-value=0.05) signify statistically significant negative correlations. (E-F) Spearman correlation analysis of log10(transactivation activity) of 27 IDRs-SOX2 versus their S ratio (in Panel E) or EDKRH ratio (in Panel F) in 27 IDRs. The red or green line shows the best-fit regression; shaded indicates the 95% confidence interval. (G) Five IDRs, rich in S but lacking EDKRH, whose condensates are miscible with Pol II condensates, were selected for fusion with the charge-rich peptide to increase charge levels. Matrices display their condensates as miscible (soft pink) and immiscible (light gray) when paired with Pol II condensates. (H) Results of dual-luciferase assays to assess the transcriptional activity of artificial TFs. Each IDR was tested with or without the charge-rich peptide (Charge). (I) A disorder prediction for human native full-length SOX2 by IUPred2A (https://iupred2a.elte.hu/). (J) Ratios of EDKRH and S in the IDRs of SOX2wt, SOX2mut1 (substituting EDKRH with S), and SOX2mut2 (substituting S with EDKRH). (K) Fluorescence imaging of condensates formed by SOX2(IDR)wt, SOX2(IDR)mut1, and SOX2(IDR)mut2 in cells, facilitated by Optodroplet system. (L-M) Fluorescence imaging and quantitative co-localization analysis of Pol II condensates with condensates formed by SOX2(IDR)wt, SOX2(IDR)mut1, and SOX2(IDR)mut2 in cells. (N) qRT-PCR analysis was conducted to measure the relative expression levels of SOX2 target genes, including *VRK1*, *ABCC2*, and *F2R*, activated by different forms of SOX2 (SOX2(IDR)wt, SOX2(IDR)mut1, and SOX2(IDR)mut2), with GAPDH serving as the internal reference.

Next, we investigated the relationship between amino acid patterns and the transactivation activity of the IDRs-SOX2 TFs. The results indicated that the ratio of EDKRH and positively charged AAs (NCPR value, and the ratios of R, K, and KRH) negatively correlated with transactivation activity (p = 0.027), while the ratio of S displayed a moderately positive, though marginally significant, correlation (p = 0.070) (Figure 7D-F). Similar patterns were observed when analyzing the relationship between the ratio of S or EDKRH in IDRs and their condensate miscibility with Pol II condensates (Figure S7A-C). Subsequently, we selected the five IDRs enriched in S and devoid of EDKRH, whose condensates are miscible with Pol II condensates, and fused them with a charge-rich peptide (named Charge) to examine how additional charge affects their miscibility and transactivation activity. This modification led to three of the condensates losing their ability to mix with Pol II condensates (Figures 7G, S7D). The dual-luciferase assay revealed that four of the tested Charge-IDRs-SOX2 variants, including the two that were miscible with Pol II condensates, exhibited reduced transactivation activity, with Charge-FUS-SOX2 as an exception (Figure 7H, Table S5). These findings suggest that a high ratio of charges may disrupt miscibility with Pol II condensates and decrease transactivation activity.

In subsequent analysis, we explored whether similar behaviors occurred in native TFs. We selected TFs^84^ containing long IDRs and analyzed their activation and repression domains. Our analysis revealed that repression domains had significantly higher ratios of charged amino acids compared to activation domains (Figure S7E, Table S6). These findings were consistent with our observed correlations between AA patterns and the transactivation activity (Figure 7D). Additionally, we also found that activation domains exhibited significantly higher ratios of G and Q (Figure S7E), which warrant further investigation.

We then investigated the impact of altering the ratio of S or charged amino acids in the native TFs, including SOX2, OCT4, and N-myc. Disorder prediction analysis indicated that the C-terminal region of full-length human SOX2 is disordered (Figure 7I). All wild-type and mutated SOX2 IDRs, including mut1 (substituting charged amino acids with S) and mut2 (substituting S with charged amino acids), could form condensates in cells (Figure 7J-K, Document S1). Notably, the condensates formed by SOX2(IDR)wt and SOX2(IDR)mut1 exhibited high miscibility with Pol II condensates, whereas SOX2(IDR)mut2 showed significantly reduced miscibility (Figure 7L-M). To assess the impact of these substitutions on the transactivation activity of SOX2, we used qRT-PCR to measure the expression levels of *VRK1^85^*, *ABCC2^86^*, and *F2R^87^*, all activated by SOX2 and containing multiple SOX2 binding motifs in their promoters (Figure 7N). The results demonstrated that SOX2mut1 significantly enhanced transactivation activity, whereas SOX2mut2 showed a marked decrease (Figure 7N). Similar patterns were also observed in OCT4 and N-myc, both known to undergo phase separation^88,89^ driven by IDRs (Figure S7F, S7L). Substituting charged AAs with S in these TFs (Figure S7G, S7M, and Document S1) maintained the ability to undergo phase separation (Figure S7H, S7N), maintained or improved miscibility with Pol II condensates (Figure S7I-J, S7O-P), and enhanced transactivation activity, as evidenced by elevated expression of target genes^89-94^ (Figure S7K, S7Q). Conversely, substituting S with charged AAs significantly reduced their miscibility (Figure S7I-J, S7O-P) and transactivation activity (Figure S7K, S7Q).

In conclusion, the miscibility between TFs and Pol II condensates influences the strength of transcription activity: high ratios of charged AAs in the IDRs of TFs affect their miscibility with Pol II condensates and downregulate their transactivation activity.

## Discussion

Over the past decade, extensive research has been conducted to unravel the mechanisms behind condensate formation and their selective partitioning or exclusion; however, regarding the coexistence of condensates, only a few case-based studies exist, highlighting a notable lack of common rules. This paper comprehensively distills the rules governing the coexistence of different condensates (Figure S6D). We found that, out of the 378 pairwise combinations of 28 different IDRs, 100 exhibited miscibility while 278 demonstrated immiscibility. Most of these findings were novel, with only a handful (5.6%, 21/378) having been previously studied^27,29-33^ (Document S2). 18/21 pairs were consistent with our findings. Different from our findings, Gilat et al. reported that FUS_IDR_/TDP43_IDR_, hnRNPA1_IDR_/TDP43_IDR_, and DDX4_IDR_/TDP43_IDR_, were immiscible^30^. In their study, the formation of persistent condensates was facilitated by constitutive oligomers; furthermore, a short region within TDP43_IDR_ that forms an α-helix^30,95^ significantly enhances homotypic interactions^96-99^, while over time, TDP43_IDR_ condensates tend to solidify^99^, collectively contributing to its condensate immiscibility. In contrast, our experiments employed the Optodroplet and Corelets systems, inducing rapid condensate formation that was imaged within minutes, a duration too short for hardening. Therefore, when studying the coexistence behavior of different condensates, the use of light-induced systems may be an ideal strategy.

We found that serine generally enhances condensate miscibility, particularly for aromatic-rich condensates. The increased miscibility facilitated by serine may be due to its potential to weaken homotypic interactions, thus lowering the barrier to mixing. Recent studies reported that a high ratio of serine corresponds to a higher effective solvation volume^6^, which represents the average space allocated per residue for solvent interactions^100-102^, and IDRs with a higher effective solvation volume tend to discourage the formation of homotypic interactions^71,72^. In this study, we increased the serine content of IDRs by fusing them with a serine-rich peptide at the terminus. Our results suggest that this modification was an effective way to enhance condensate miscibility. Further experiments are necessary to fully determine whether the miscibility-enhancing role of serine is influenced by its distribution within the protein.

Many proteins rich in oppositely charged AAs—including DDX4^13^, LAF-1^12,56,103^, NPM1^104^, CBX2^105^, Tau^106,107^, and FMRP^61^—have been found to undergo phase separation driven by homotypic electrostatic interactions. Our findings suggest that IDRs with opposite charges prefer forming homotypic interactions that drive the formation of condensates, and disfavor forming heterotypic interactions, thereby inhibiting the mixing between charge-rich condensates. When these charged amino acids cluster into blocks, heterotypic interactions are significantly altered, with charge-blocky condensates tending toward miscibility. This behavior has been corroborated by several independent studies^32,35,108^. Moreover, in this study, we selected IDRs lacking charges and fused them with a charge-dispersed peptide, enriched with alternating positive and negative charges. Our results suggest that this alternation was an effective way to promote condensate immiscibility. Further experiments are required to ascertain the differences between the effects of dispersed charges, achieved through alternating positive and negative charges, versus those achieved by randomly distributed charges.

Many post-translational modifications (PTMs) can significantly alter the characteristics of proteins^109-118^, which further affect in condensates formation^8,15,57,61,64,119-129^, selective partitioning^31,130^, and the coexistence behavior of different condensates^131^. For instance, phosphorylation of the linker protein SAPAP enhances the mixing of PSD-95-positive condensates with Shank3-positive condensates by improving the PPIs^131^. In our study, we found that over 70% of disordered proteins (8672 out of 12111) could undergo phosphorylation, affecting their phase separation capabilities and potentially altering condensate miscibility. Focusing on SRSF1, which exhibits the highest phosphorylation levels of serine and tyrosine, we noted that while SRSF1 condensates can mix with Pol II condensates, phosphorylation of SRSF1 impedes this miscibility. This observation suggests that the dynamic regulation of condensate mixing or demixing through specific PTMs may be a common cellular strategy.

Over the past decade, numerous transcriptional condensates have been identified, both in physiology^32,52,132-134^ and in diseases^34,55,89,135-137^. These condensates regulate transcription by partitioning coactivators and Pol II. In this study, we found that the miscibility between TF condensates and Pol II condensates positively correlates with transactivation activity, reinforcing the critical role of phase separation in transcriptional regulation^33,68,138^. However, we also observed that TFs exhibiting high miscibility with Pol II condensates do not invariably lead to gene activation, suggesting that additional regulators might need to be co-compartmentalized to enhance transcription. Moreover, TFs exhibiting low miscibility with Pol II do not consistently repress gene expression. This observation aligns with our previous findings^32^ where condensates formed by TFs, such as PITX2 or ZIC3, do not mix with Pol II condensates, yet free Pol II can still moderately enter PITX2 or ZIC3 condensates and activate gene expression, albeit at a reduced level. We propose that some specific transcriptional condensates, even when only slightly partitioning free Pol II, can still function effectively. Future research should aim to more clearly distinguish the roles of partitioning and mixing in transcription regulation. Lastly, it is important to note that all findings in this study regarding transcriptional condensates were contextual, limited to promoter regions containing multiple binding motif repeats, artificial TFs with long IDRs driving phase separation, and overexpression to reach the concentration thresholds necessary for phase separation. Therefore, we must acknowledge that while phase separation can regulate transcription, it is not universally necessary or sufficient for the process.

## Limitation

This study has illuminated key mechanisms by which specific amino acid compositions influence the miscibility of condensates. However, several limitations warrant further investigation. Among our tested IDRs, Y served as the primary aromatic amino acid, without F and W. It remains to be seen whether the roles of aromatic amino acids in affecting condensate miscibility would be consistent if F or W were the primary aromatic contributors. Additionally, in our examination of charged amino acids, we did not differentiate among K, R, and H, nor between E and D. Our analysis did not include polyQ, G, P, and hydrophobic AAs, which may also contribute to condensate formation. Lastly, as our experiments were conducted in cellular environments, it is crucial to consider that these condensates partition many native proteins or even RNAs, potentially affecting their properties.

## Acknowledgments

We thank Benjamin R. Sabari, Bing Li, Cong Liu, Heankel Lyons, Weijie Chen, Yi Lin, and Yinqing Li for helpful comments and discussion. This work was supported by grants from the the Natural Science Foundation of China (32125010 and 32330024 to P.L.; T2325003 and 32070666 to T.L.), Beijing Natural Science Foundation of (Z230014 to P.L. and T.L.) and National Key R&D Program (2023YFF1204703 to P.L. and T.L.; 2021YFF1200900 to T.L.). G.P. was supported by the Natural Science Foundation of China (32400560), the Postdoctoral Fellowship Program of CPSF under Grant Number GZC20240859, the Shuimu Tsinghua Scholar Program (2023SM250), the Postdoctoral Program at the Tsinghua University–Peking University Joint Center for Life Sciences, and the Advanced Innovation Fellow Program. We are grateful to the Nikon Biological Imaging Center (Tsinghua University, Beijing, China) for confocal imaging support.

## Author Contributions

Conceptualization, G.P., T.L., and P.L.; Methodology, G.P., and X.W.; Investigation, G.P., X.W., D.G., W.X., and Z.C.; Writing-original draft, G.P.; Writing-review&editing, T.L., and P.L.; Funding acquisition, P.L., T.L., and G.P.

## STAR METHODS

### RESOURCE AVAILABILITY

#### Lead contact

Further information and requests for resources and reagents should be directed to and will be fulfilled by the Lead Contact Pilong Li.

#### Materials availability

The study did not generate new unique reagents.

#### Data and code availability

##### Data

All data reported in this paper will be shared by the lead contact upon request.

##### Code

All original code is available in this paper’s supplemental information.

Any additional information required to reanalyze the data reported in this paper is available from the lead contact upon request

#### Experimental model and study participant details

##### Cell culture

HEK293T cells were cultured in Dulbecco’s modified Eagle’s medium (HyClone, Inc.) supplemented with 10% fetal bovine serum (Invitrogen, Inc.) at 37℃ with 5% CO2. FuGENE HD (Promega, Inc.) was used for transient transfection according to the manufacturer’s instructions.

### METHOD DETAILS

#### Cloning

In this study, we utilized various truncated, full-length, or mutated proteins as follows: BRD4^52^ (Aa 673-1350), BRD9^68^ (Aa 3-133), CPEB2^60^ (Aa 2-138), DDX4^14^ (Aa 1-207), FMRP^7,61^ (Aa 466-632), FUS^62^ (Aa 1-214), HES4^68^ (Aa 1-68), hnRNPA1^63^ (Aa 186-320), hnRNPA2^64^ (Aa 193-353), hnRNPA3^10^ (Aa 207-378), hnRNPAB^10^ (Aa 235-332), hnRNPD^10^ (Aa 262-355), hnRNPDL^10^ (Aa 316-420), hnRNPH1^10^ (Aa 365-452), hnRNPK^65^ (Aa 1-463), MED1^52^ (Aa 948-1575), MED15^68^ (Aa 1-424), MLLT1^68^ (Aa 186-484), NUP98^54^ (Aa 1-500), Pol II^66^ (Aa 1593-1960), PRCC^68^ (Aa 1-156), RBM14^67^ (Aa 257-573), SFPQ^68^ (Aa 265-565), SMARCA2^68^ (Aa 1-254), SOX2 DBD (Aa 39-121), SRSF1 (Aa 1-248), SRSF1-5StoD (Aa 1-248), SRSF1-10StoD (Aa 1-248), SRSF1-20StoD (Aa 1-248), CLK1 (Aa 1-484), TAF15(IDR1)^10,69^ (Aa 1-156), TAF15(IDR2)^69^ (Aa 417-512), TDP43^10^ (Aa 258-414), TIA1^70^ (Aa 291-386), SOX2 (Aa 1-317), SOX2(IDR) (Aa 122-317), OCT4 (Aa 1-360), OCT4(IDR) (Aa 1-136, 289-360), N-myc (Aa 1-464), N-myc(IDR) (Aa 1-392) (Table S1). All proteins, with the exception of BRD4, MED1, and Pol II, which are derived from mice, were human in origin. The genes for these proteins were synthesized by Genewiz and subsequently the corresponding fragments were amplified and cloned into either the pcDNA3.1 or pETL7 plasmid vectors, contingent on the specific requirements of our experiments.

#### Calculation of NCPR and Hydropathy

Features of NCPR and Hydropathy^139^ are calculated by localCIDER 0.1.21^78,140^, using functions of localCIDER.SequenceParameters().get_NCPR() and localCIDER.SequenceParameters().get_mean_hydropathy(). (Table S1)

#### Analyzing the Miscibility Index and Amino Acid Patterns

Calculate Miscibility Index: For each of the 28 IDRs, calculate the Miscibility Index by averaging the Pearson correlation coefficients obtained from comparing the condensate of the specific IDR with the condensates of each of the other 27 IDRs individually.

Rank IDRs: Rank all 28 IDRs based on their Miscibility Index. (Table S2)

Divide into Groups: For each cutoff value from 4 to 24, divide the IDRs into two groups: The top N (4 to 24) IDRs are classified as the ‘High-miscible Group’. The remaining IDRs (24 to 4) are classified as the ‘Low-miscible Group’.

Assess Differences in Amino Acid Patterns: For each division of IDRs into high and low miscible groups, assess the statistical significance of the differences in 26 amino acid patterns between the two groups.

Generate P-values: This process generates multiple sets of p-values (21 sets, corresponding to each cutoff from 4 to 24) for each amino acid pattern.

Determine Significance: The final significance of each amino acid pattern is determined by averaging the five lowest p-values from the 21 sets generated. (Figure 1F-G)

#### Blocky Charge Distribution Analysis

First, we devised an analytical framework where every 10-amino-acid segment was employed as a sliding window to quantify charge blocks. We defined segments with net charge per residue (NCPR) values of ≥0.5 as positive and those with values of ≤-0.5 as negative, adjacent blocks were subsequently merged. We identified four as ‘charge-blocky’, characterized by having at least three charge blocks, with at least one being of opposite charge. Additionally, we identified seven as ‘charge-dispersed’, five of which contained only one charge block; notably, two IDRs without charge blocks were also categorized as ‘charge-dispersed’ due to their high ratio of charged AAs (Figure S5C)

#### Identifying IDR-Containing Proteins

We acquired annotations of disordered regions for the human proteome from the PhaSePred website, saved in the Human_reviewed.json file. These disordered regions include low-complexity domains as predicted by SEG, prion-like domains by PLAAC, and intrinsically disordered domains by Espritz. We also retrieved structural data for human proteins from the AlphaFold2 database, specifically from the file UP000005640_9606_HUMAN_v4.tar. Regions with a pLDDT score below 50 were classified as disordered. If the sequences between two adjacent disordered regions were less than 10 amino acids, these regions were merged into a single disordered region. Among these, 18,985 disordered regions exceed 50 amino acids in length and are distributed across 12,111 proteins. (Figure 2F, Table S3)

#### Acquisition of Weak PPIN

We retrieved the human interactome dataset (BIOGRID-ORGANISM-Homo_sapiens-4.4.230.tab3.txt) from the BioGRID database. This dataset was narrowed down to interactions where both interactors, labeled as Organism Name Interactor A and Organism Name Interactor B, were identified as Homo sapiens. We further refined these interactions based on the type of Experimental System used. Additionally, from a total of 12,111 IDR-containing proteins, we isolated those PPIs documented by Proximity-Labeling Mass Spectrometry (PL-MS), specifically excluding those identified by Yeast Two-Hybrid (Y2H) and Affinity Capture-MS (AC-MS). This selection process helped us pinpoint a subset of proteins that form weak PPIN. Ultimately, we identified 51,152 weak PPIs involving 5,651 proteins. (Figure 2F, Table S3)

#### PPI Densities Analysis

To compare the PPI densities between S(Rich) and S(Poor) groups, we first ranked these proteins within weak PPIN according to their S ratio. We then selected the top 200, 250, 300, 350, 400, 450, and 500 proteins to form seven S(Rich) groups, and the bottom 200, 250, 300, 350, 400, 450, and 500 to form seven S(Poor) groups. We calculated the PPI densities for each group and subsequently performed statistical analysis to compare the PPI densities between the S(Rich) groups and the S(Poor) groups (Figure 2G).

To compare the PPI densities between FYW(Rich) and FYW(Poor) groups, we first ranked these proteins within weak PPIN according to their FYW ratio. We then selected the top 200, 250, 300, 350, 400, 450, and 500 proteins to form seven FYW(Rich) groups, and the bottom 200, 250, 300, 350, 400, 450, and 500 to form seven FYW(Poor) groups. We calculated the PPI densities for each group using the Python package NetworkX 3.1. By extracting the subnetworks formed within each group, we applied the density function from NetworkX to compute the connectivity density. The algorithm is as follows: Density=2M/(N(N-1)), where M is the number of edges in the subnetwork, and N is the number of nodes in the network. Finally, we performed statistical analysis to compare the PPI densities between the FYW(Rich) groups and the FYW(Poor) groups (Figure 3D).

To compare the PPI densities between FYW(Rich)&S(Rich) groups and FYW(Rich)&S(Poor) groups, we first ranked these proteins within weak PPIN according to their FYW ratio, selecting the top 1,000 as FYW(Rich) proteins. We then re-ranked the FYW(Rich) proteins by their S ratio, selecting the top 200, 250, 300, 350, 400, 450, and 500 to form seven FYW(Rich)&S(Rich) groups, and the bottom 200, 250, 300, 350, 400, 450, and 500 to form seven FYW(Rich)&S(Poor) groups. Finally, we calculated the PPI densities for each group and performed a statistical analysis to compare the PPI densities between FYW(Rich)&S(Rich) groups and FYW(Rich)&S(Poor) groups (Figure 3I).

To compare PPI densities between two groups—(positive NCPR paired with negative NCPR) and (positive NCPR paired with neutral NCPR)—we first ranked these proteins within weak PPIN according to their NCPR values. We selected the top 200, 250, 300, 350, 400, 450, and 500 proteins to form seven positive NCPR groups, and the bottom 200, 250, 300, 350, 400, 450, and 500 to form seven negative NCPR groups. Similarly, we selected proteins nearest to zero NCPR in increments of 200, 250, 300, 350, 400, 450, and 500 to establish seven neutral NCPR groups. For each group pairing, the top 200 proteins from the positive NCPR groups were paired with the corresponding bottom 200 from the negative NCPR groups, and with the 200 proteins closest to zero from the neutral NCPR groups, for which we calculated PPI densities. This process was repeated using six additional cutoffs: 250, 300, 350, 400, 450, and 500. Finally, we conducted statistical analyses to compare the PPI densities between these group pairings (Figure 4E).

To compare the PPI densities between two groups—(ED(High) paired with KRH(High)) and (ED(High) paired with KRH(Low))—we first ranked these proteins within weak PPIN according to their ED ratio and selected the top 200, 250, 300, 350, 400, 450, and 500 to form seven ED(High) groups. We then re-ranked the remaining proteins according to their KRH ratio, selecting the top and bottom 200, 250, 300, 350, 400, 450, and 500 to form seven KRH(High) and KRH(Low) groups. We used the top 200 of the ED(High) groups to pair with the top 200 of the KRH(High) group, and calculated the PPI densities. This process was replicated using six additional cutoffs: 250, 300, 350, 400, 450, and 500. Consequently, we obtained seven PPI densities for the groups of (ED(High) paired with KRH(High)). Similar procedures were performed to obtain seven PPI densities for the group of (ED(High) paired with KRH(Low)). Finally, we performed statistical analysis to compare the PPI densities between these two groupings (Figure 4F).

To compare the PPI densities in EDKRH(High) groups with those in EDKRH(Low) groups, we ranked these proteins within weak PPIN according to their EDKRH ratio, selecting the top 200, 250, 300, 350, 400, 450, and 500 to form seven EDKRH(High) groups, and the bottom 200, 250, 300, 350, 400, 450, and 500, creating seven EDKRH(Low) groups. Finally, we calculated the PPI densities for each group and performed a statistical analysis to compare the PPI densities between the EDKRH(High) and EDKRH(Low) groups (Figure 4J).

To compare PPI densities in three group pairings—((EDKRH(High) paired with FYW(Rich)), and (EDKRH(Low) paired with FYW(Rich))—we began by ranking these proteins within weak PPIN according to their FYW ratio. We selected the top 200, 250, 300, 350, 400, 450, and 500 proteins to form seven FYW(Rich) groups. Subsequently, we re-ranked the remaining proteins according to their ED ratio, selecting the top 1,000 as ED(High) proteins and the bottom 1,000 as ED(Low) proteins. The ED(High) proteins were then re-ranked by their KRH ratio and selected the top 200, 250, 300, 350, 400, 450, and 500 proteins to form seven EDKRH(High) groups. Similarly, the ED(Low) proteins were re-ranked by their KRH ratio, with the bottom 200, 250, 300, 350, 400, 450, and 500 forming seven EDKRH(Low) groups. For each group pairing, the top 200 proteins from the FYW(Rich) groups were paired with the corresponding top 200 from the EDKRH(High) groups and bottom 200 from the EDKRH(Low) groups, for which PPI densities were calculated. This procedure was replicated across six additional cutoffs: 250, 300, 350, 400, 450, and 500. Finally, we performed statistical analyses to compare the PPI densities across the two group pairings (Figure 4N).

To compare the PPI densities among EDKRH(High)&S(Poor) groups, EDKRH(High)&S(Rich) groups and EDKRH(Low)&S(Rich) groups, we first ranked these proteins within weak PPIN according to their EDKRH ratio, selecting the top 1,000 as EDKRH(High) proteins and the bottom 1,000 as EDKRH(Low) proteins. We re-ranked the EDKRH(Low) proteins by their S ratio, selecting the top 200, 250, 300, 350, 400, 450, and 500 to form seven EDKRH(Low)&S(Rich) groups. We then re-ranked the EDKRH(High) proteins by their S ratio, selecting the top 200, 250, 300, 350, 400, 450, and 500 to form seven EDKRH(High)&S(Rich) groups, and the bottom 200, 250, 300, 350, 400, 450, and 500 to form seven EDKRH(High)&S(Poor) groups. Finally, we calculated the PPI densities for each group and performed a statistical analysis to compare the PPI densities among EDKRH(High)&S(Poor) groups, EDKRH(High)&S(Rich) groups and EDKRH(Low)&S(Rich) groups (Figure 5E).

#### Phosphorylation Level Analysis

Human phosphorylation data were sourced from the PhosphoSite database (version 2023.12). We identified phosphorylated sites on IDRs and calculated the phosphorylation ratios for serine and tyrosine (S+Y). The analysis revealed 8,672 proteins encompassing 12,888 IDRs with at least one phosphorylated S/Y site. Among these, 2,945 proteins, each with a SaPS score of ≥ 0.5, contained a total of 5,322 IDRs. (Table S4)

#### Phase Separation Prediction

PS(Reported): A total of 703 human phase-separating (PS) proteins were identified from 1,419 entries in PhaSepDB2.1^141^, based on low-throughput experimental evidence, along with 770 entries related to membraneless organelles. Additionally, 85 human PS proteins were retrieved from LLPSDB^142^, and 59 from the PhaSePro^143^ database. The union of these datasets resulted in 725 known human PS proteins. Of these, 582 proteins contain disordered regions, which we selected for further analysis, designated as PS(Reported).

PS(Predicted) and NoPS(Predicted): Proteins predicted to undergo phase separation (PS) or not (NoPS) were identified from the PhaSePred database (http://predict.phasep.pro/download/), using a SaPS score threshold of ≥ 0.5 for PS(Predicted) and < 0.5 for NoPS(Predicted), excluding those already present in the known human PS protein set.

The serine/tyrosine phosphorylation ratios of these proteins were then compared using the Mann-Whitney U-test from the Python package scipy (version 1.11.4). (Table S4)

#### Gene Ontology (GO) Enrichment Analysis

We ranked 5,322 IDRs from 2,945 proteins, each containing at least one S/Y phosphorylation site and having a SaPS score of ≥ 0.5. We then selected the top 20% of IDRs, resulting in a dataset of 1,064 IDRs from 822 proteins. Gene Ontology (GO) analysis was performed on these 822 proteins using the online tool DAVID^144^, with the background set to all 12,111 human disordered proteins.

#### Mutagenesis of Native Transcription Factors

For SOX2mut1, its IDR contains only a few charged amino acids; therefore, in addition to these, selected glycine and alanine residues were also substituted with serine. For OCT4mut1 and N-mycmut1, the modifications primarily involved replacing the charged amino acids within the IDRs with serine. In the case of SOX2mut2, OCT4mut2, and N-mycmut2, the mutations predominantly consisted of substituting serine residues in the IDRs with charged amino acids.

#### Sequence Alignment

The sequence alignments were performed using the online tool available at MultAlin^145^ (http://multalin.toulouse.inra.fr/multalin/multalin.html). We utilized the default parameters set by the website for all alignments. The results of these alignments can be found in Document S1.

#### Protein Purification

The recombinant plasmids encoding MBP-SRSF1-His, MBP-SRSF1-5StoD-His, MBP-SRSF1-10StoD-His, MBP-SRSF1-20StoD-His, MBP-CLK1-His, and MBP-Pol His were successfully transformed into E. coli BL21 (DE3) cells. Isolated single-cell clones were cultivated in LB broth at 37°C until the culture’s optical density at 600 nm reached 0.8. Protein overexpression was then induced with 0.5 mM IPTG and allowed to proceed at 16°C overnight. Post-induction, cells were harvested, resuspended in a lysis buffer containing 40 mM Tris-HCl and 500 mM NaCl at pH 8.0, and subsequently disrupted by sonication. Cell debris was removed by centrifugation at 15,000 rpm for 30 minutes. The supernatant containing His-tagged proteins was then subjected to purification using Ni-NTA agarose (Thermo Fisher Scientific, Catalog #R90115), followed by ion-exchange and gel-filtration chromatography steps for further purification. Only high-quality proteins, as confirmed by SDS-PAGE, were concentrated using Amicon Ultra-4 spin columns (Merck-Millipore). These proteins were then stored at -80°C in a buffer consisting of 20 mM HEPES, 500 mM NaCl, and 10% [v/v] glycerol, pH 7.4, to ensure their stability for future use.

#### Protein Labeling

The purified proteins were labeled with Alexa Fluor 488 or 546 (Thermo Fisher Scientific), with the reaction conducted at room temperature for one hour to ensure thorough binding of the dye. Following this incubation, the illustra™ Microspin G-50 Columns (GE Healthcare) were employed to effectively remove any excess dye not bound to the proteins.

#### Protein Condensate Assays

For the condensate assays in vitro, protein samples were diluted to the desired concentrations in a buffer that resulted in a final composition of 20 mM HEPES (pH 7.4), 150 mM NaCl, and 10% (v/v) glycerol within low-binding microscopy plates (Cellvis, P384-1.5H-N). Concurrently, TEV protease was added to the same plate to facilitate the cleavage of the MBP tag. Following a one-hour incubation at room temperature, the samples were examined for phase separation.

#### Microscopy

Imaging was with Nikon A1R HD25 Laser Scanning Confocal Microscope. The resulting data were analyzed using the NIS-Elements AR software, version 5.2.

#### Live-cell Imaging

Cells were grown in 24-well glass bottom plates (Cellvis, P24-1.5H-N). Live-cell imaging was performed using a Nikon A1R HD25 Laser Scanning Confocal Microscope. This setup included a temperature-controlled incubation chamber maintained at 37℃ with 5% CO2. For activating the cells, we used specific laser wavelengths: 488 nm both for Cry2 and for iLID/SspB activation, as well as for imaging the GFP. Additionally, a 560 nm wavelength was employed for imaging mCherry. The duration of light stimulation ranged from 30 to 60 seconds, and the duration of imaging was in minutes, which was sufficient to allow IDR condensation, yet short enough to prevent potential hardening of the condensates.

#### Luciferase Assay

A promoter with 26 repeats of the CATTGTT sequence, recognized by SOX2(DBD), was inserted into the pGL4 plasmid before the firefly luciferase gene. These repeats were separated by random nucleotides ranging from 5 to 20 bases. HEK 293T cells were separately co-transfected with 27 artificial TFs IDRs-SOX2 and the 26× CATTGTT-contained firefly luciferase plasmid, along with a Renilla luciferase plasmid (Promega, E6921) to serve as a reference for transfection efficiency. Post-transfection (approximately 24 hours), dual-luciferase assays were employed to quantify reporter gene activity, as per the manufacturer’s protocol provided with the Promega Dual-Luciferase Reporter Assay System (catalog number E1960). Quantification of firefly luciferase emission was normalized to Renilla luciferase signal to account for variability in transfection efficiency. This normalized value was then compared to that of a control vector expressing SOX2(DBD). To ensure reproducibility and statistical validity, each experiment was three times. (Table S5)

#### Amino Acid Analysis of TFs

We obtained transcription activation and repression domains from the published paper^84^. We selected sequences annotated as having at least 75% disordered regions. We then calculated and compared the amino acid characteristics between the transcription activation and repression domains. We used the stats.mannwhitneyu() function from the Python scipy package to perform a U-test and calculate the p-value. (Table S6)

#### Phosphorylation Reactions

Phosphorylation assays were performed using 50 μM of purified MBP-SRSF1-His as substrate, with 50 nM of purified CLK1 kinase and 200 μM ATP in 1× kinase buffer (Cell Signaling Technology, 9802). The mixtures were incubated at 37°C, and aliquots were taken at 0, 20, 40, and 60 minutes to terminate the reactions by adding EDTA to a final concentration of 5 mM. The phosphorylation levels of SRSF1 were assessed via Western blotting.

#### Western Blotting

Proteins underwent fractionation through SDS-PAGE and were then transferred onto a PVDF membrane. This membrane was incubated overnight with a rabbit polyclonal antibody specific to SRSF1 (Beyotime, AF8058). Subsequently, an HRP-conjugated goat anti-rabbit secondary antibody (Thermo Fisher Scientific, A24537) was applied. For visualization, we used an enhanced chemiluminescence reagent.

#### Fisher’s Exact Test

Performed Fisher’s exact test calculations using the stats.fisher_exact() function from the Python package scipy version 1.10.1.

### KEY RESOURCES TABLE

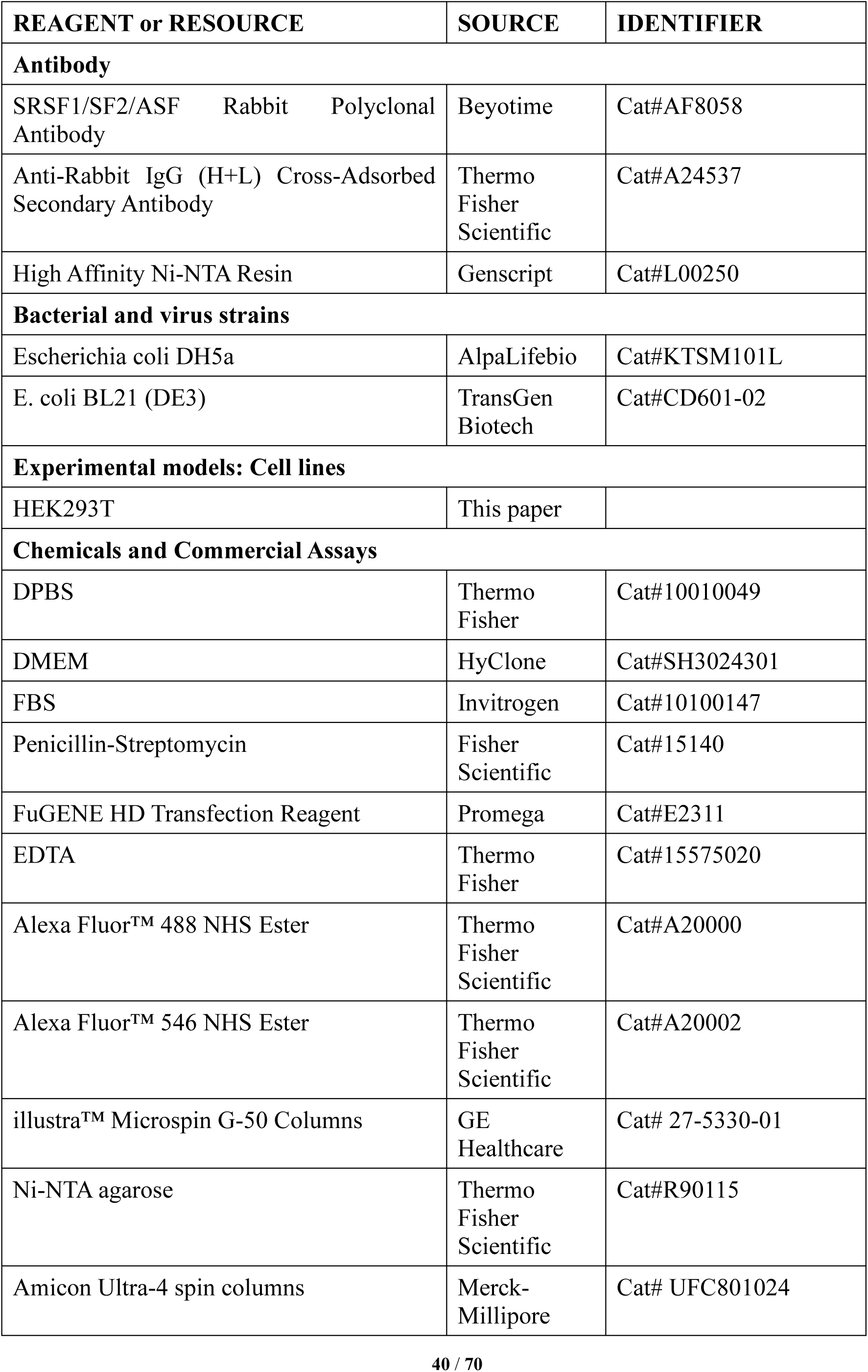
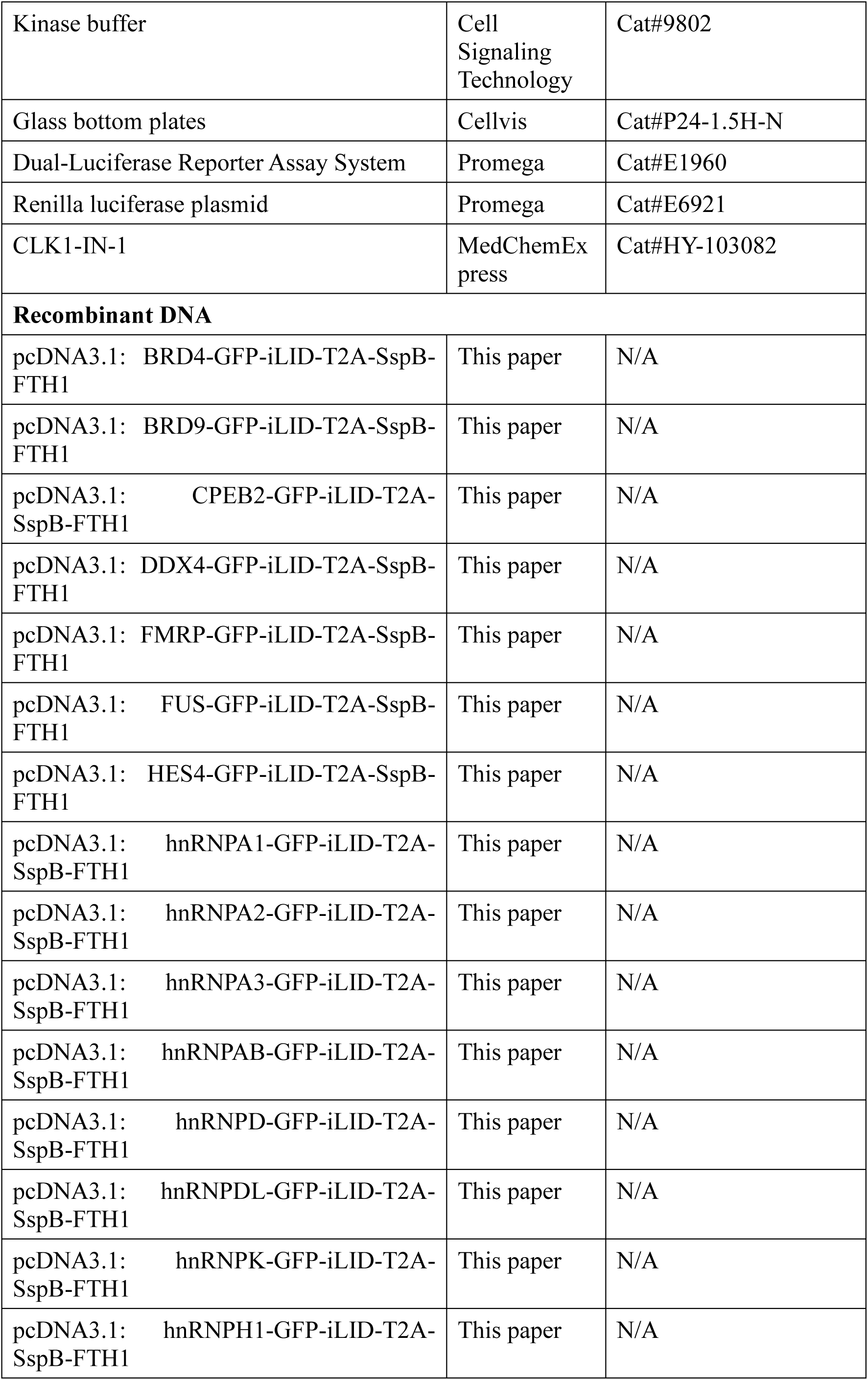
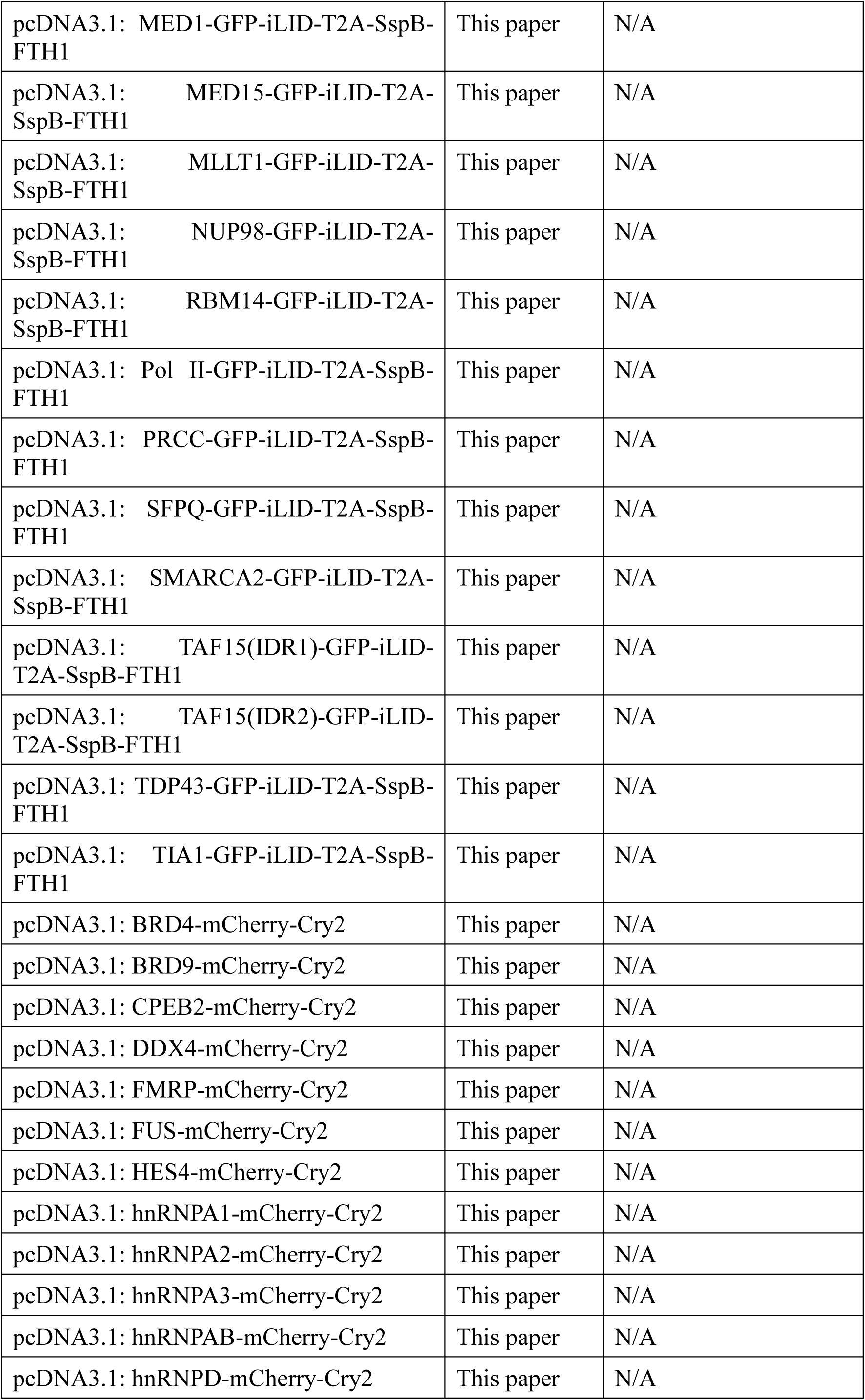
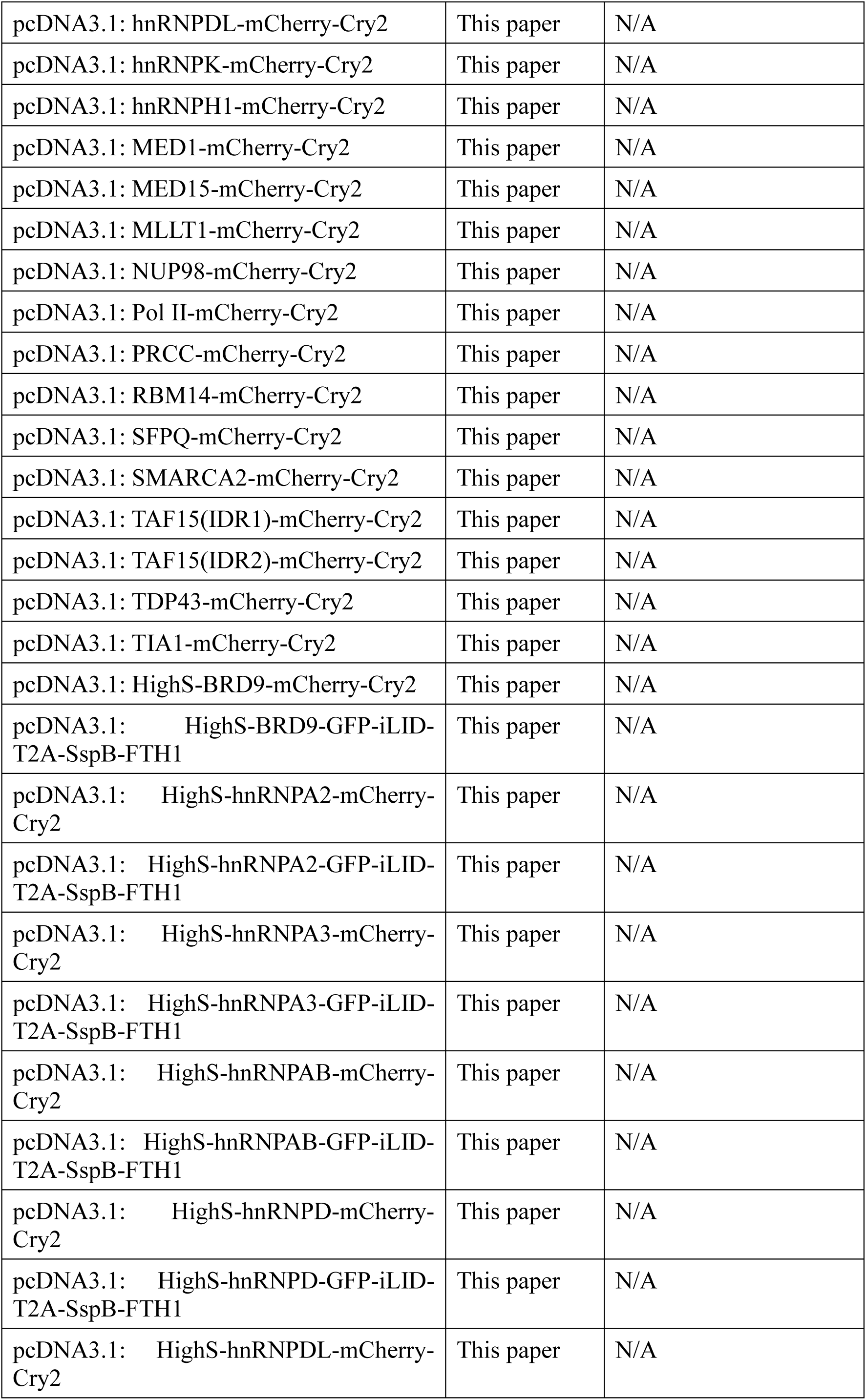
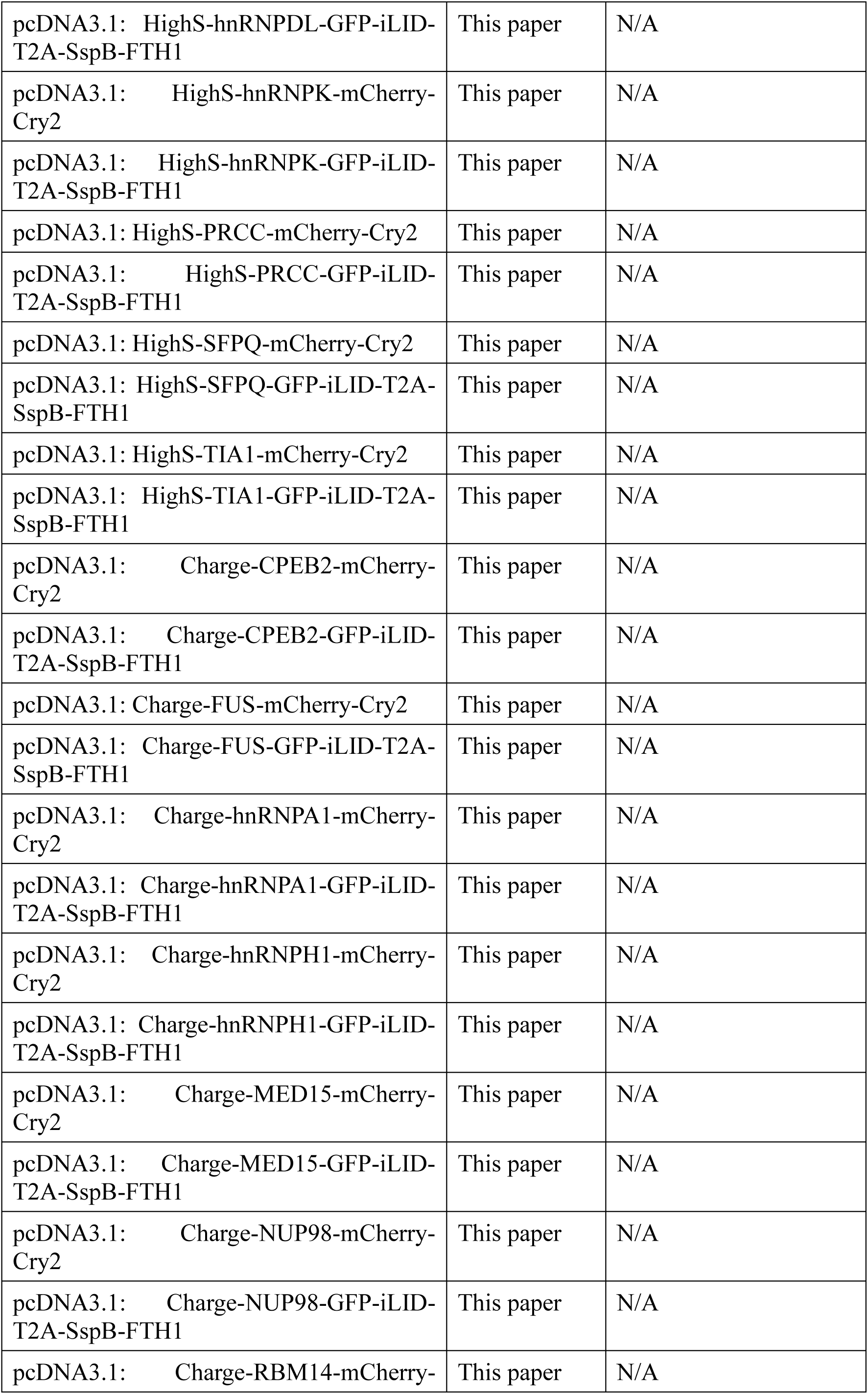
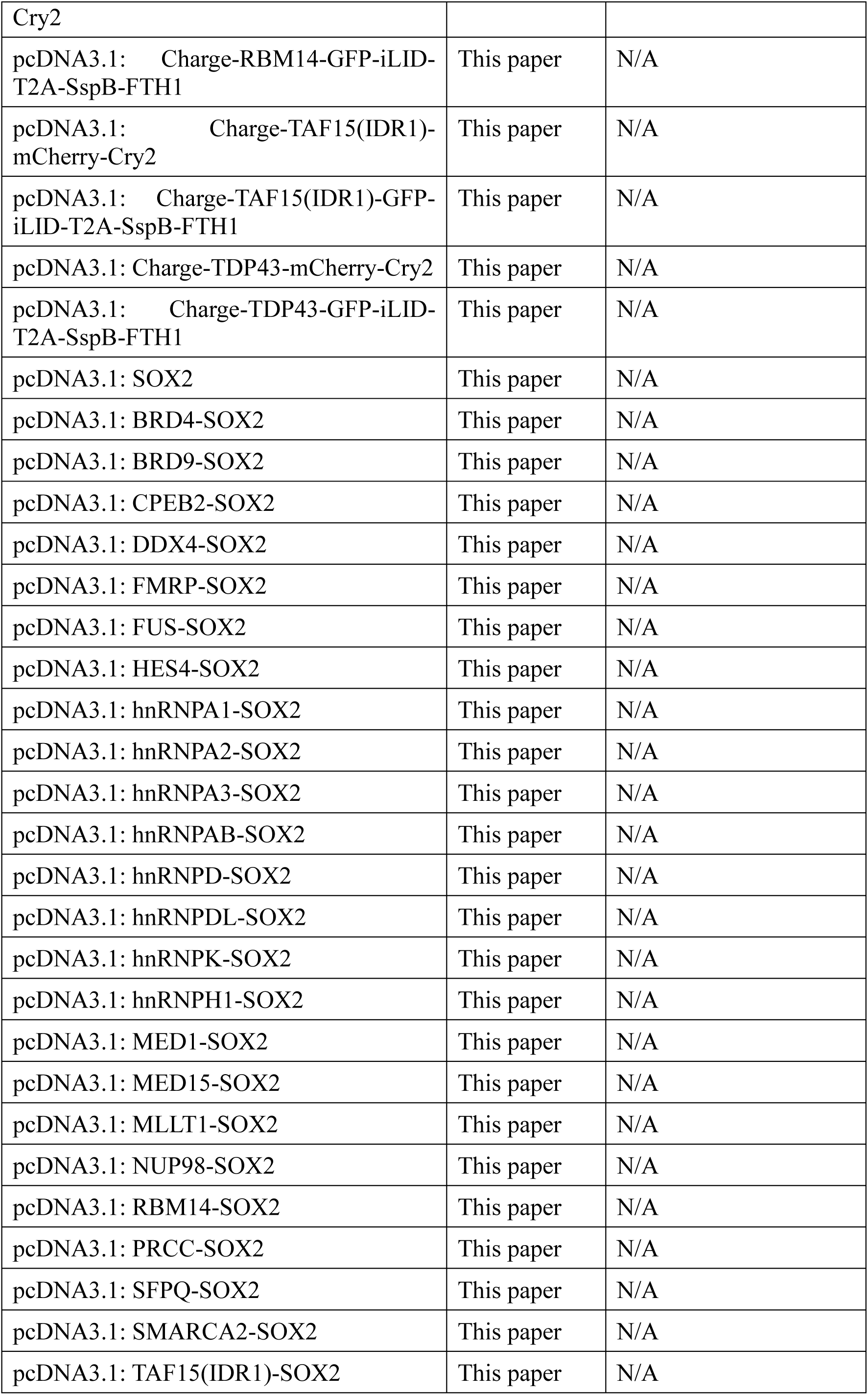
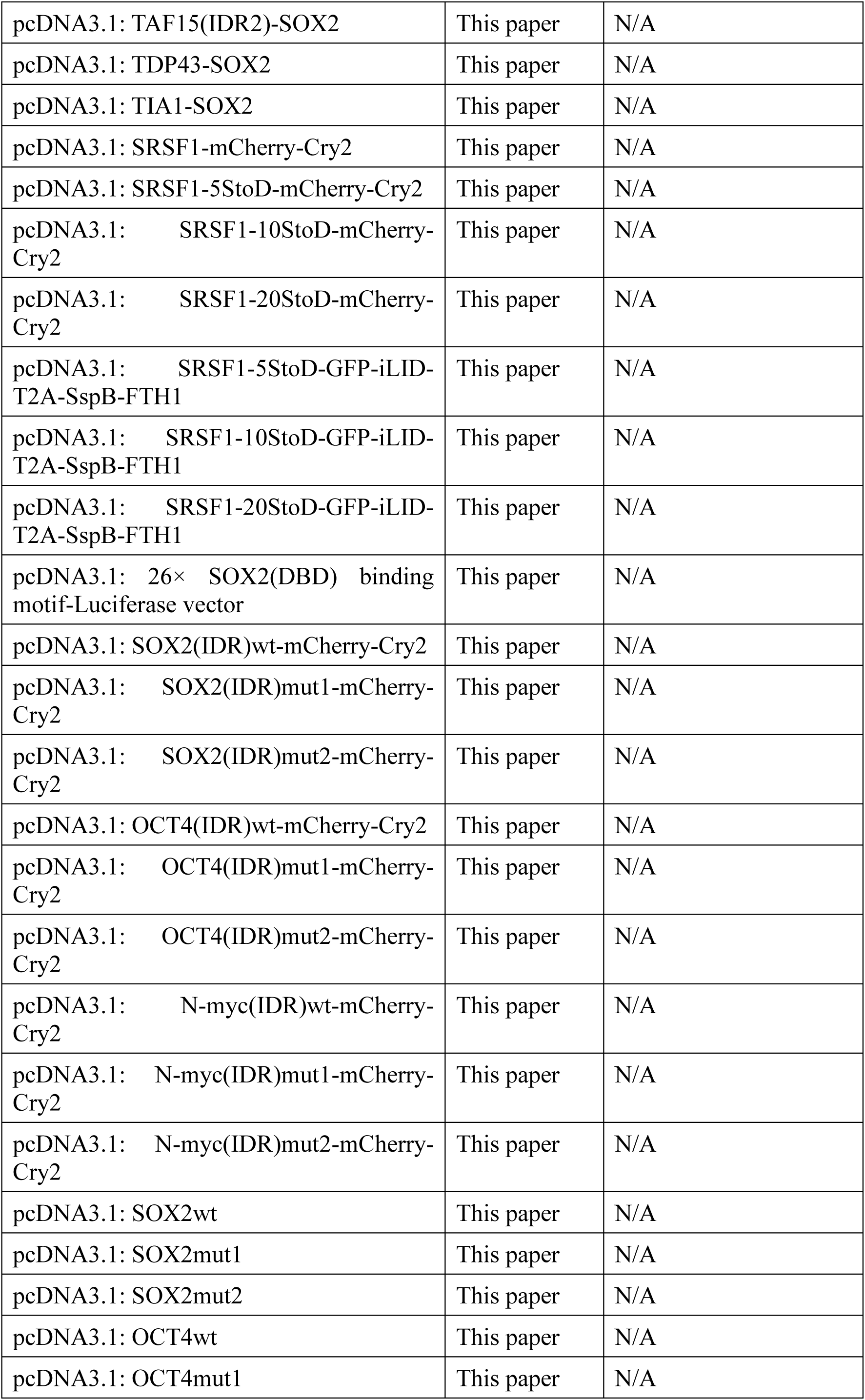
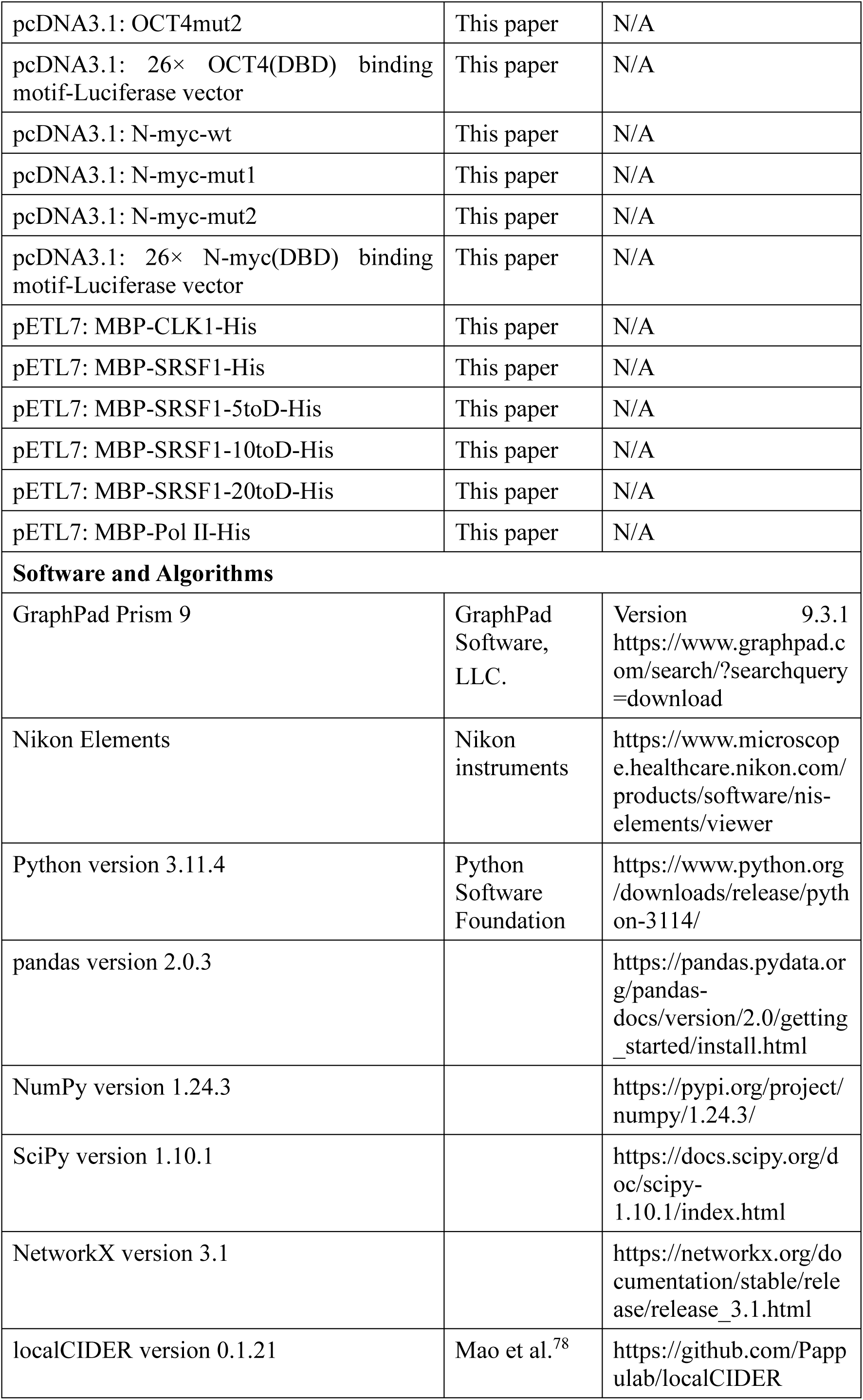

## Supplemental Figures Legends

**Figure S1.**
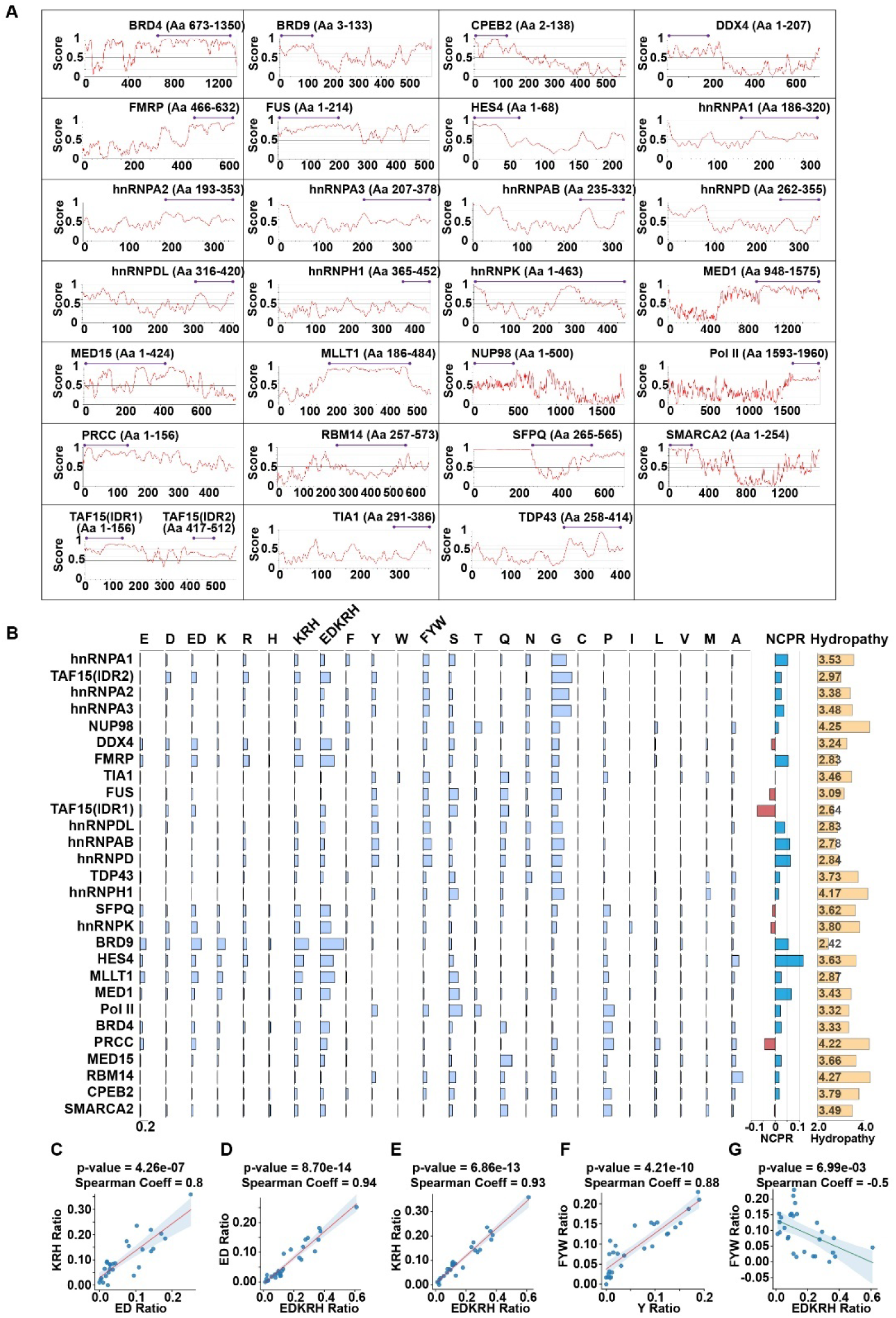
Amino Acid Composition and Properties of Selected IDRs. (A) Selection of 28 intrinsic disordered regions (IDRs) from 27 proteins for analysis. TAF15 contains two IDRs. IDRs are predicted by IUPred2A (https://iupred2a.elte.hu/). (B) Chart presenting the amino acid (AA) composition and properties of 28 IDRs. Light-blue boxes indicate the ratio of each AA in the IDRs. The scale bar at the bottom left shows a standard ratio of 0.2. Positive and negative NCPR (net charge per residue) values are presented as blue and red bars, respectively. Hydropathy values are indicated at the right. (C) Spearman correlation analysis for ED ratio versus KRH ratio across the 28 IDRs. The red line represents the best-fit regression line, and the shaded area depicts the 95% confidence interval. (D) Spearman correlation analysis for EDKRH ratio versus ED ratio across the 28 IDRs. Data are presented as in Panel C. (E) Spearman correlation analysis for EDKRH ratio versus KRH ratio across the 28 IDRs. Data are presented as in Panel C. (F) Spearman correlation analysis for Y ratio versus FYW ratio across the 28 IDRs. Data are presented as in Panel C. (G) Spearman correlation analysis for EDKRH ratio versus FYW ratio across the 28 IDRs. Data are presented as in Panel C.

**Figure S2.**
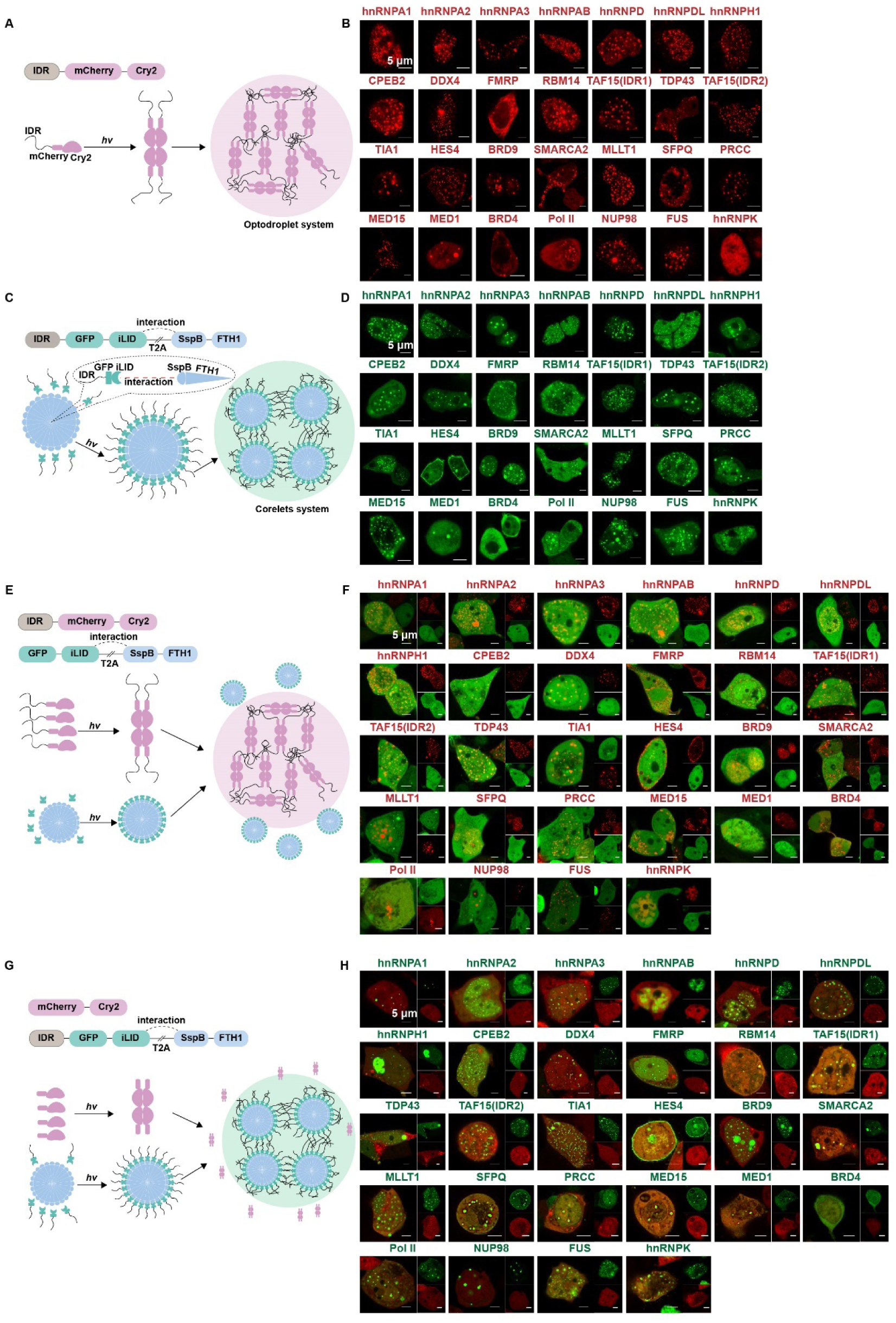
Light-induced Phase Separation Mechanisms. (A) Schematic representation of the phase separation process for IDRs fused with mCherry-Cry2 upon blue light exposure. Theoretically, exposing cells overexpressing IDRs-mCherry-Cry2 constructs to blue light causes Cry2 to oligomerize, which amplifies multivalent interactions within the IDR and induces phase separation (B) Cellular fluorescence microscopy shows IDRs-mCherry-Cry2 undergoing phase separation upon blue light induction. (C) Schematic representation of the phase separation process for IDRs fused with GFP-iLID-T2A-SspB-FTH1 upon blue light exposure. Theoretically, in cells overexpressing these constructs, the T2A peptide crucially enabled ribosomal peptide bond skipping, yielding two distinct proteins: IDRs-GFP-iLID and SspB-FTH1. FTH1 naturally forms a homo-24mer structure. Therefore, under blue light exposure, iLID interacted with SspB, facilitating the formation of a heteromeric 24-mer complex between IDRs-GFP-iLID and SspB-FTH1, which then undergo phase separation. (D) Cellular fluorescence microscopy shows IDRs-GFP-iLID-T2A-SspB-FTH1 undergoing phase separation upon blue light induction. (E) Schematic representation shows GFP-iLID/SspB-FTH1 was excluded by IDRs-mCherry-Cry2 condensates. (F) Fluorescence images demonstrate the specific exclusion of GFP-iLID/SspB-FTH1 from IDRs-mCherry-Cry2 condensates. (G) Schematic representation shows mCherry-Cry2 was excluded by IDRs-GFP-iLID/SspB-FTH1 condensates. (H) Fluorescence images demonstrate the specific exclusion of mCherry-Cry2 from IDRs-GFP-iLID/SspB-FTH1 condensates.

**Figure S3.**
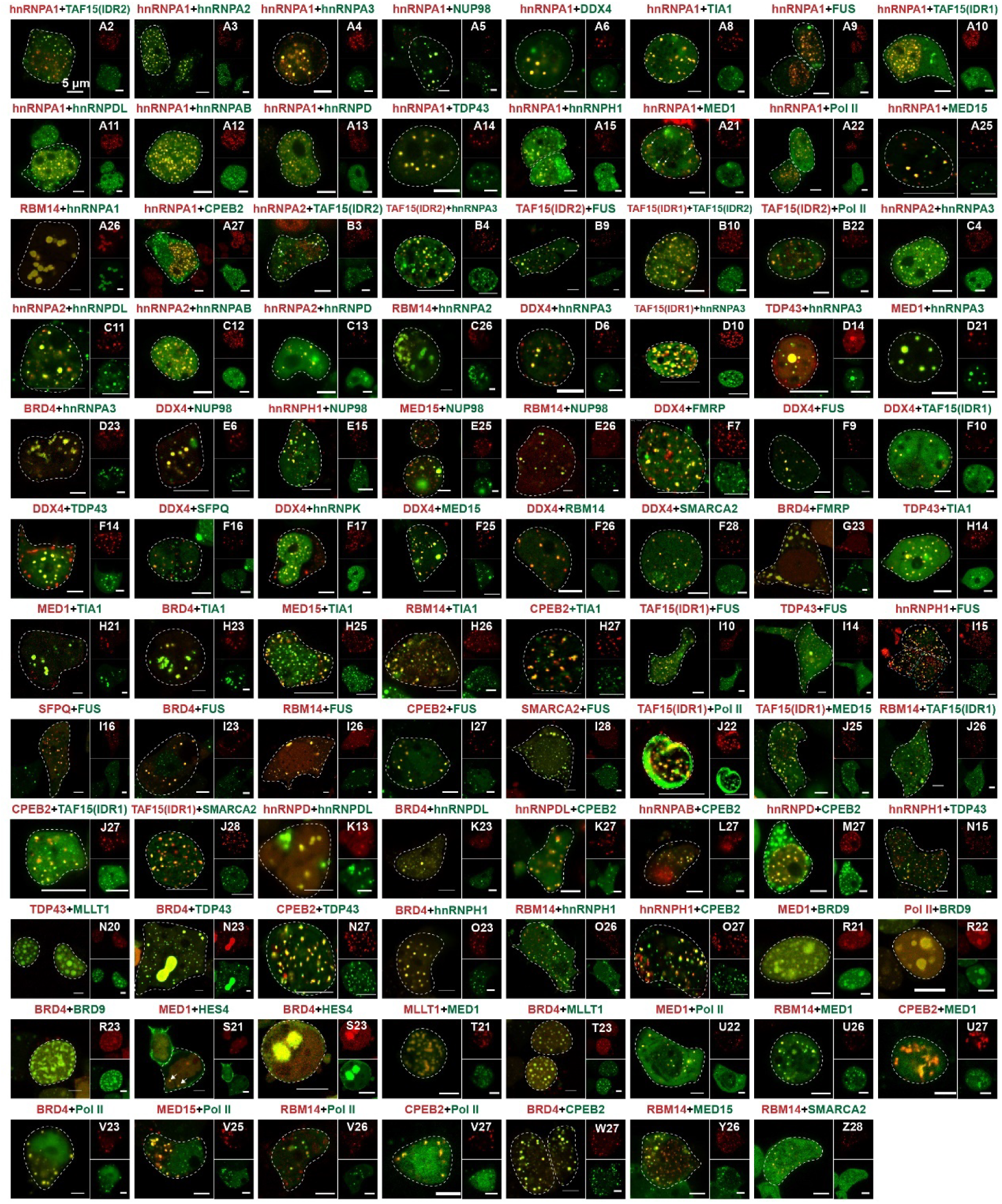
Fluorescence Imaging of Miscible Condensates. Related to Figure 1. This figure presents the fluorescence microscopy images of all miscible condensate pairs formed by 28 IDRs. The labels at the top-right of each image correspond to the row and column from Figure 1E.

**Figure S4.**
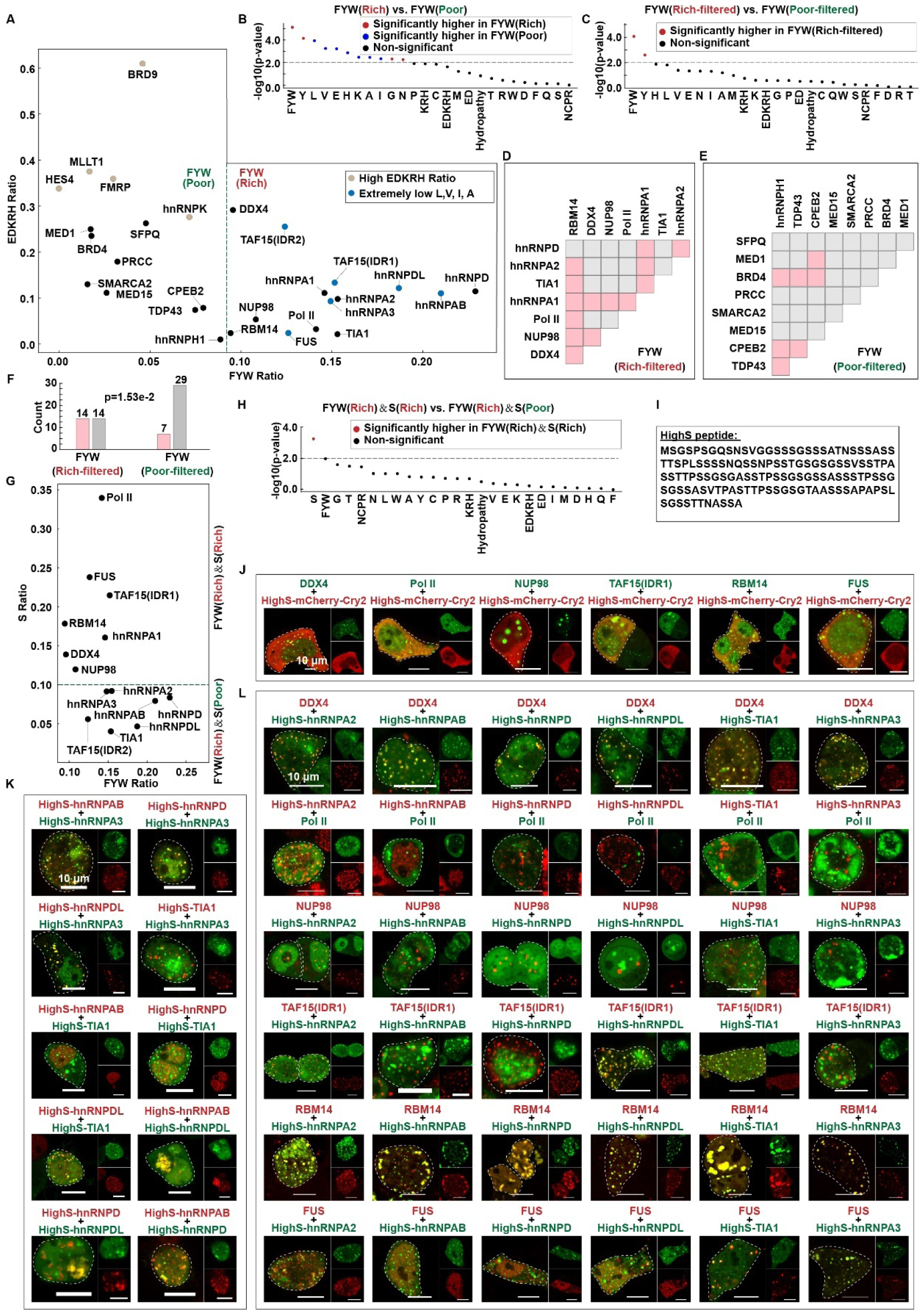
Additional Serine Enhances Miscibility of Aromatic-rich IDR Condensates. Related to Figure 3. (A) Scatter plot displaying the distribution of IDRs based on their FYW and EDKRH ratios. The IDRs are categorized into two main groups: FYW(Rich) and FYW(Poor). IDRs in the FYW(Rich) group, marked in deep blue, are excluded due to extremely low ratios of the amino acids L, V, I, and A; the remaining IDRs are FYW(Poor) group, marked in gray, are excluded due to high EDKRH ratios; the remaining IDRs are referred to as FYW(Poor-filtered) and are displayed in Panel E. (B) Plot showing statistical differences in the 26 AA patterns between IDRs categorized as FYW(Rich) and those categorized as FYW(Poor). Red dots indicate patterns significantly more prevalent in the FYW(Rich) group, whereas blue dots indicate patterns significantly more prevalent in the FYW(Poor) group. Black dots represent non-significant differences. (C) Plot showing statistical differences in the 26 AA patterns between IDRs in the FYW(Rich-filtered) group and the FYW(Poor-filtered) group. Red dots indicate patterns significantly enriched in the filtered FYW(Rich-filtered) group. Black dots signify non-significant differences. (D) Matrix showing miscible (soft pink) and immiscible (light gray) pairs of condensates formed by FYW(Rich-filtered) IDRs. (E) Matrix showing miscible (soft pink) and immiscible (light gray) pairs of condensates formed by FYW(Poor-filtered) IDRs. (F) Quantification of the data from Panels D and E. Fisher’s exact test was performed. (G) Scatter plot showing the distribution of FYW-rich IDRs (from Figure 3A) based on their S and FYW ratios. IDRs are categorized into two groups: FYW(Rich)&S(Rich) and FYW(Rich)&S(Poor). (H) Plot presenting statistical differences in the 26 AA patterns between IDRs in the FYW(Rich)&S(Rich) group and IDRs in the FYW(Rich)&S(Poor) group. Red dots indicate patterns significantly enriched in the FYW(Rich)&S(Rich) group. Black dots signify non-significant differences. (I) The sequence of the serine-rich peptide (HighS). (J) Fluorescence images showing that the HighS-mCherry-Cry2 fusion protein does not form condensates and does not partition into condensates formed by FYW(Rich)&HighS IDRs. (K) Fluorescence images of miscible and immiscible pairs of condensates formed by FYW(Rich)&HighS IDRs. (L) Fluorescence images of miscible and immiscible condensates formed by FYW(Rich)&S(Rich) IDRs paired with condensates formed by FYW(Rich)&HighS IDRs.

**Figure S5.**
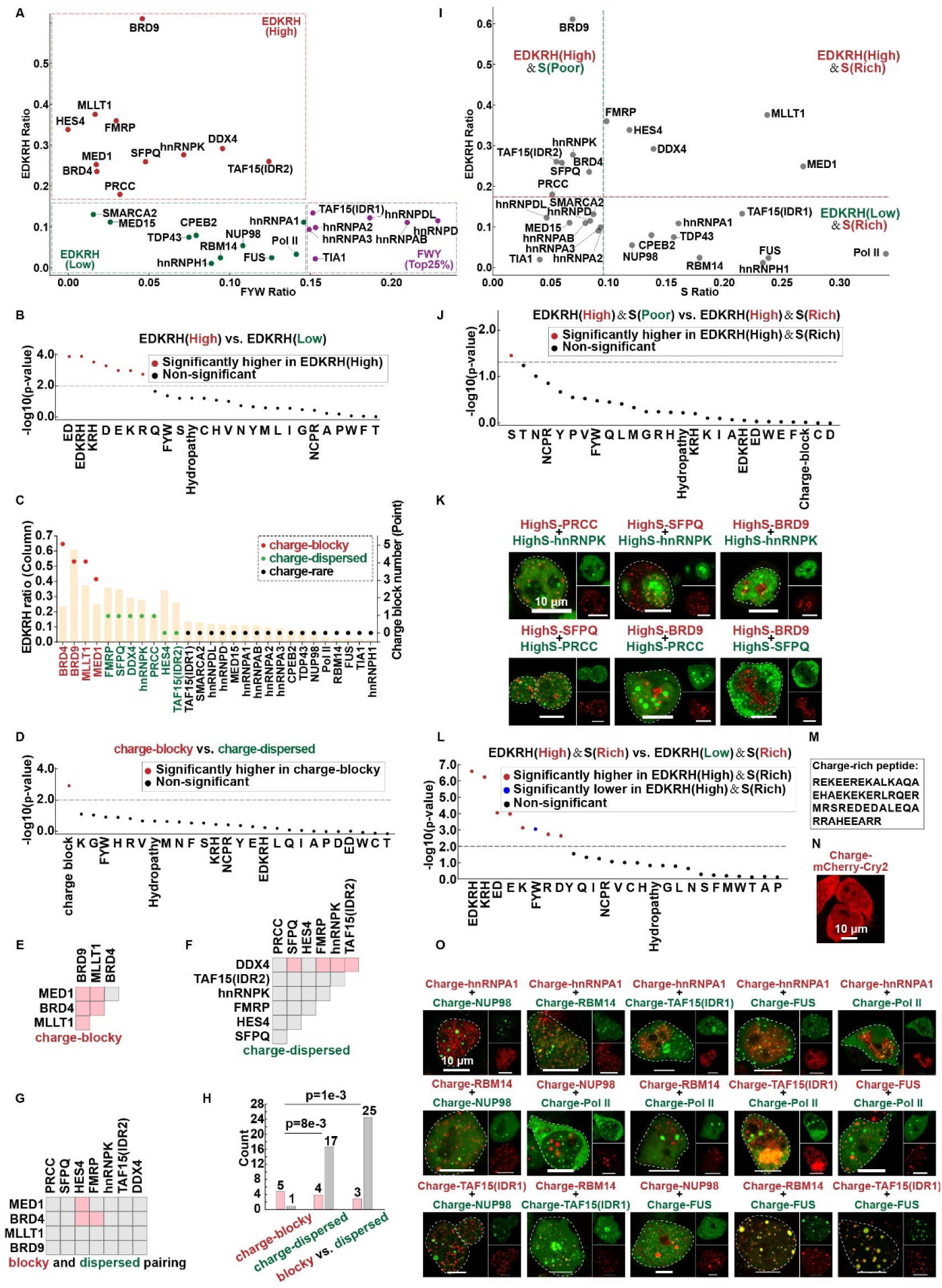
Charge Promotes Condensate Immiscibility. Related to Figure 4 and Figure 5. (A) Scatter plot displaying the distribution of IDRs based on their FYW and EDKRH ratios. The top 25% of IDRs with the highest FYW ratios were excluded, and the remaining IDRs were categorized into two groups: EDKRH(High) and EDKRH(Low). (B) Plot showing statistical differences in the AA patterns between IDRs categorized as EDKRH(High) and IDRs categorized as EDKRH(Low). Red dots indicate patterns significantly enriched in the EDKRH(High) group. Black dots represent non-significant differences. (C) Chart displaying the ratios of charged amino acids (columns) and counts of charge blocks (points) across 28 IDRs. Charge-blocky, charge-dispersed, and charge-rare IDRs are marked in red, green, and black, respectively. (D) Plot showing statistical differences in the AA patterns between IDRs categorized as ‘charge-blocky’ and IDRs categorized as ‘charge-dispersed’. Red dots indicate patterns significantly more prevalent in the charge-blocky group. Black dots represent non-significant differences. (E-G) Matrices showing miscibility (soft pink) and immiscibility (light gray) of condensates: (E) formed by pairs of charge-blocky IDRs, (F) formed by pairs of charge-dispersed IDRs, and (G) formed by charge-blocky IDRs paired with charge-dispersed IDRs. (H) Quantification of the data from Panels E-G. Fisher’s exact test was performed (I) Scatter plot displaying the distribution of IDRs based on their S and EDKRH ratios. IDRs were categorized into four groups: EDKRH(High)&S(Rich), EDKRH(High)&S(Poor), EDKRH(Low)&S(Rich), and EDKRH(Low)&S(Poor). (J) Plot showing statistical differences in the AA patterns between IDRs categorized as EDKRH(High)&S(Rich) and IDRs categorized as EDKRH(High)&S(Poor). Red dots indicate patterns significantly enriched in the EDKRH(High)&S(Rich) group. Black dots represent non-significant differences. (K) Fluorescence images of miscible and immiscible pairs of condensates formed by EDKRH(High)&HighS IDRs. (L) Plot showing statistical differences in the AA patterns between IDRs categorized as EDKRH(High)&S(Rich) and IDRs categorized as EDKRH(Low)&S(Rich). Red dots indicate patterns significantly more prevalent in the EDKRH(High)&S(Rich) group. Blue dots indicate patterns significantly less prevalent in the EDKRH(High)&S(Rich) group. Black dots represent non-significant differences. (M) The sequence of the charge-rich peptide (Charge). (N) Fluorescence images demonstrating that the charge-rich peptide fused with mCherry-Cry2 does not form condensates upon light induction. (O) Fluorescence images of miscible and immiscible pairs of condensates formed by High-EDKRH(High)&S(Rich) IDRs.

**Figure S6.**
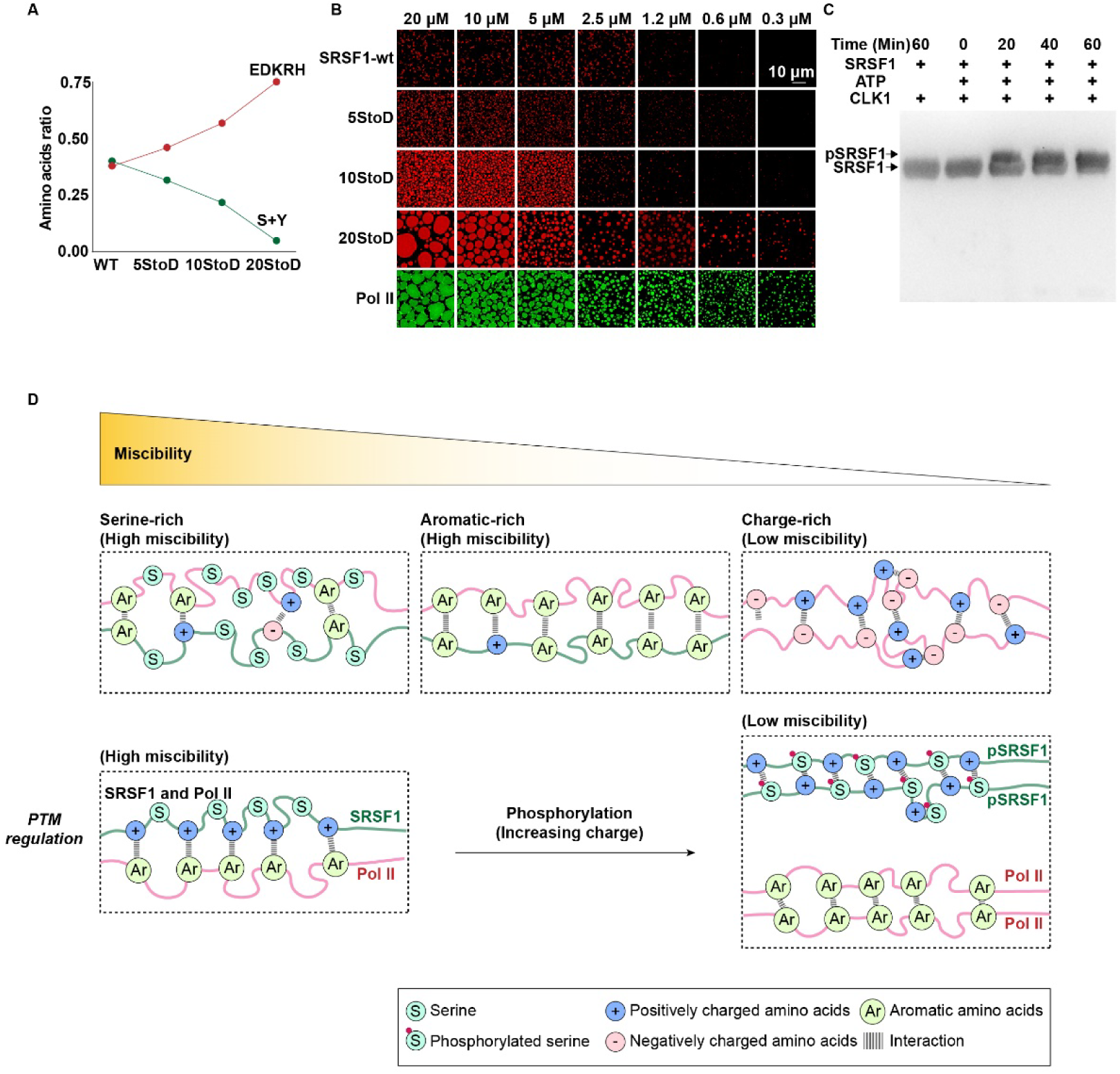
Phosphorylation-enhanced Charges in SRSF1 Reduce Miscibility with Pol II Condensates. Related to Figure 6. (A) Ratios of EDKRH and S+Y amino acids in the RS domain of SRSF1 wildtype and variants. (B) Fluorescence images displaying condensates formed by the full-length wild-type SRSF1, its variants, and the wild-type Pol II IDR in vitro. (C) Western blot showing SRSF1 phosphorylation levels modified by CLK1. (D) Schematic diagram illustrating the condensate miscibility. Condensates enriched in serine or aromatic amino acids display high miscibility. Condensates characterized by high levels of charge show very low miscibility. Post-translational modifications (PTMs) can dynamically alter the miscibility of these condensates. For instance, SRSF1 condensates initially mix well with Pol II condensates; however, phosphorylation of SRSF1, which introduces charges, reduces this miscibility.

**Figure S7.**
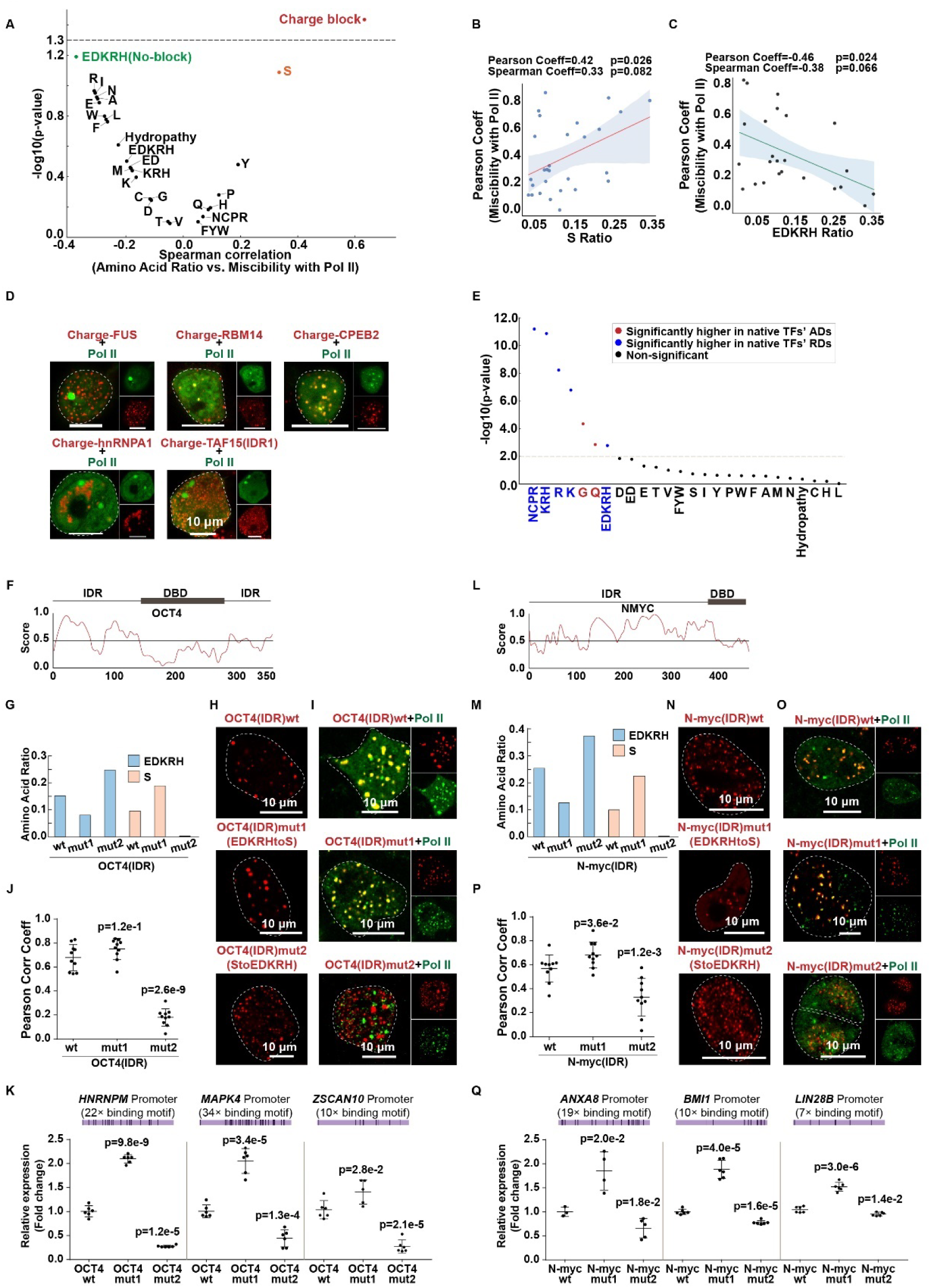
Highly Charged TFs Condensates Show Low Miscibility with Pol II Condensates and Decreased Transactivation. Related to Figure 7. (A) Volcano plot illustrating the Spearman correlation analysis between amino acid patterns in 27 IDRs and their miscibility with Pol II condensates. The x-axis shows the Spearman correlation coefficient, and the y-axis the negative log10 of the p-values. (B) Spearman and Pearson correlation analysis of the S ratio in IDRs versus their miscibility with Pol II condensates. Pearson coefficient is 0.42 (p=0.026); Spearman coefficient is 0.33 (p=0.082). The red line indicates the regression best-fit, with a shaded area for the 95% confidence interval. (C) Spearman and Pearson correlation analysis of the EDKRH ratio in 24 IDRs (excluding four with charge blocks) versus their miscibility with Pol II condensates. Pearson coefficient is -0.46 (p=0.024); Spearman coefficient is -0.38 (p=0.066). The green line indicates the best-fit regression; shading indicates the 95% confidence interval. (D) Fluorescence images of miscible and immiscible condensates formed by Charge-IDRs paired with condensates formed by Pol II. (E) Plot showing statistical differences in the 26 AA patterns between 212 activation domains (ADs) and 93 repression domains (RDs) from 234 native TFs. (F) A disorder prediction for human native full-length OCT4 by IUPred2A (https://iupred2a.elte.hu/). (G) Ratios of EDKRH and S in the IDRs of OCT4wt, OCT4mut1 (substituting EDKRH with S), and OCT4mut2 (substituting S with EDKRH). (H) Fluorescence imaging of condensates formed by OCT4(IDR)wt, OCT4(IDR)mut1, and OCT4(IDR)mut2 in cells, facilitated by Optodroplet system. (I-J) Fluorescence imaging and quantitative co-localization analysis of Pol II condensates with condensates formed by OCT4(IDR)wt, OCT4(IDR)mut1, and OCT4(IDR)mut2 in cells. (K) qRT-PCR analysis was conducted to measure the relative expression levels of OCT4 target genes, including *HNRNPM*^90^, *MAPK4*^91^, and *ZSCAN10*^92^, activated by different forms of OCT4 (OCT4wt, OCT4mut1, and OCT4mut2), with GAPDH serving as the internal reference. (L) A disorder prediction for human native full-length N-myc by IUPred2A (https://iupred2a.elte.hu/). (M) Ratios of EDKRH and S in the IDRs of N-myc-wt, N-myc-mut1 (substituting EDKRH with S), and N-myc-mut2 (substituting S with EDKRH). (N) Fluorescence imaging of condensates formed by N-myc(IDR)wt, N-myc(IDR)mut1, and N-myc(IDR)mut2 in cells, facilitated by Optodroplet system. (O-P) Fluorescence imaging and quantitative co-localization analysis of Pol II condensates with condensates formed by N-myc(IDR)wt, N-myc(IDR)mut1, and N-myc(IDR)mut2 in cells. (Q) qRT-PCR analysis was conducted to measure the relative expression levels of N-myc target genes, including *ANXA8*^89^, *BMI1*^93^, and *LIN28B*^94^, activated by different forms of N-myc (N-myc-wt, N-myc-mut1, and N-myc-mut2), with GAPDH serving as the internal reference.

## Supplemental Document and Tables Legends

**Document S1. Sequence alignment**

**Document S2. Condensates combinations have been reported**

**Table S1. Amino acid composition analysis of 28 IDRs**

**Sheet1.** Amino acid composition analysis of 28 intrinsically disordered proteins.

**Table S2. Condensates miscibility analysis**

**Sheet1.** Assessing miscibility among 378 pairwise combinations of 28 IDRs using Pearson correlation coefficients.

**Sheet2.** Assessing miscibility index of 28 IDRs by averaging the Pearson correlation coefficients with other 27 IDR condensates.

**Table S3. PPIN analysis**

**Sheet1.** 18985-IDRs-in-12111-proteins: 18,985 IDRs exceeding 50 amino acids across 12,111 proteins.

**Sheet2.** 51152weakPPIs-in-5651-proteins: 51,152 weak protein-protein interactions involving 5,651 proteins with IDRs

**Sheet3.** 10051-IDRs-in-5651 proteins: 5651 proteins contain 10,051 IDRs.

**Sheet4.** 5651-proteins: The list of 5,651 proteins

**Sheets 5-14.** Assessing PPI densities from weak PPIN in different protein groups.

**Table S4. Phase separation prediction of phospho-proteins**

**Sheet1.** 12111-Disordered-Proteins: Utilize SEG, PLAAC, Espritz, and AlphaFold2 to identify 12,111 human proteins with disordered regions (length ≥ 50 amino acids) and employ SaPS to assess their phase separation potential in phosphorylated states.

**Sheet2.** Phosphorylation-Level-Rank: This table displays the number of proteins within various phosphorylation level percentiles that either exceed or fall below a SaPS score of 0.5, alongside the proportion of these proteins undergoing phase separation.

**Sheet3.** 2945Proteins-5322IDRs(SaPS ≥ 0.5): 2,945 phosphorylated disordered proteins show strong phase separation potential (SaPS score ≥ 0.5).

**Sheet4.** 1064IDRs-for-GO-analysis: Based on their phosphorylation levels, the top 20% of 5,322 IDRs from 2,945 proteins, each with a SaPS score ≥ 0.5, were selected for GO analysis.

**Sheet5.** Core-Splicing-Proteins: 14 core mRNA splicing proteins highlighted in Figure 6F.

**Table S5. Dual luciferase assay**

**Sheet1.** Dual-luciferase-assay-1: Use dual luciferase assay to assess the transactivation activity of 27 IDRs-SOX2 proteins.

**Sheet2.** Dual-luciferase-assay-2: Use dual luciferase assay to assess the transactivation activity of 6 charge-IDRs-SOX2 proteins.

**Table S6. Amino acid analysis of native TFs AD and RD**

**Sheet1.** TF-AD-RD-analysis: Amino acid composition analysis between the activation domain and repression domain of native transcription factors.

**Sheet2.** p-value: P-values of amino acid composition analysis between activation and repression domains.

